# Multi-cohort Analysis Reveals Microbiome Signatures Associated with Drug Response in New-Onset Rheumatoid Arthritis

**DOI:** 10.64898/2026.03.30.715090

**Authors:** Rahul L. Bodkhe, Rebecca B. Blank, Kai R. Trepka, Diego A. Orellana, Jose U. Scher, Peter J. Turnbaugh, Renuka R. Nayak

## Abstract

The human gut microbiome influences treatment outcomes, but whether microbiome signatures of drug response generalize across cohorts remains unclear. Here, we perform a multi-cohort analysis (3 cohorts, N=100 patients) to determine whether cross-cohort microbial signatures are associated with methotrexate (MTX) response in new-onset rheumatoid arthritis (RA) patients. Pre-treatment gut microbiome community structure and function differed by future MTX response status, with MTX-nonresponders (MTX-NR) showing *Bacteroides caccae* depletion and *Ruminococcus bromii* enrichment. Multiple microbial functions were depleted in MTX-NR, including nucleotide metabolism, one-carbon pool by folate, and histidine metabolism. Notably, candidate MTX-degrading genes were enriched in MTX-NR. Microbiome functional profiles outperformed clinical metrics in predicting future MTX response. These results show that consistent microbiome signatures are associated with MTX response across different RA cohorts and pave the way for microbiome-based precision medicine in newly diagnosed RA patients.

## Introduction

Our gastrointestinal tract harbors a diverse community of microorganisms, collectively known as the human gut microbiome, which plays a key role in governing individualized responses to diet^1^, pharmaceuticals^2^ and disease susceptibility^3^. Gut microbes metabolize drugs, affecting their pharmacokinetics^4,5^. Conversely, many drugs alter microbiome composition^6,7^, and in doing so, shape host immunity^7,8^. Thus, the pretreatment microbiome may be useful in predicting drug response and thereby lead to accelerated disease treatment, particularly in immune-mediated treatments. Consequently, efforts are underway to integrate these insights into precision medicine^4,9–12^.

However, a barrier to clinical translation is determining whether microbiome-drug response associations are predictive across multiple cohorts. While several studies have identified microbial taxa and functions that predict drug response^11,13–16^, it remains unclear whether microbial biomarkers of drug response are specific to a given study population or whether they generalize across cohorts. Studies of checkpoint inhibitor therapy in melanoma offer mixed findings, with some studies reporting consistent associations across cohorts whereas others report cohort-specific associations^17,18^. Here, we sought to determine whether the human gut microbiome in rheumatoid arthritis (RA) patients harbors cross-cohort microbiome signatures that predict response to methotrexate (MTX), an antimetabolite and immunomodulatory drug that interacts with the microbiome in multiple ways^4,5,7,19^. Identifying associations across cohorts would enable the development of a generalizable microbiome-based predictor that could lead to clinically actionable insights for advancing precision medicine in RA.

RA is a chronic autoimmune disease characterized by persistent inflammation of the joints, leading to pain, swelling, and progressive joint damage. MTX is widely regarded as the first-line disease-modifying anti-rheumatic drug (DMARD) due to its efficacy, cost-effectiveness, and global availability^4,20^. However, only up to 50% of patients show clinically adequate response to MTX monotherapy after a 12-week treatment period^21,22^. Given this critical “window of opportunity” to initiate effective therapy and prevent irreversible damage, there is a need for predictive biomarkers of treatment response^23^. Baseline clinical parameters^24^, pharmacogenomic variants^25^, and circulating metabolic markers, including serum lipidomic profiles, have been associated with treatment outcome^24–26^. However, most proposed predictors show limited reproducibility across independent cohorts^23,26^. These limitations highlight the need for generalizable predictive biomarkers and improved insights into mechanisms of MTX response.

Differences in MTX response may stem from inter-individual variation in gut microbiome profiles^4,9,11,27,28^. We and others have shown that the pretreatment gut microbiome community structure and function differ in patients based on their future response to MTX^4,9,11,27,28^. Furthermore, we have shown that the gut microbiome impacts MTX bioavailability and that microbial MTX metabolism is associated with differential drug response in patients^4,5,19^. Notably, both taxonomic and functional microbiome features have been linked to treatment outcomes, and predictive models based on pretreatment microbiome profiles have been explored and validated^4,9^.

However, these studies were cohort-specific, and whether microbial signatures associated with MTX response generalize across multiple cohorts remains unknown. Here, we conducted a comprehensive analysis of the human gut microbiome in new-onset RA patients by integrating datasets from three independent cohorts to (i) identify microbial features consistently associated with MTX-NR, (ii) gain mechanistic insights into MTX response, and (iii) develop a predictive model that is generalizable across populations for the early identification of MTX-NR. In the combined cohort, the pretreatment microbiome was associated with MTX-response, with taxa and functional profiles showing biologically meaningful associations with future response. Key pathways implicated in MTX biology, including nucleotide and amino acid metabolism, were depleted in MTX-NR; in contrast, candidate MTX metabolizing genes were enriched. A microbiome-based machine learning model trained on cross-cohort functional profiles accurately predicted future MTX-NR in a validation cohort. This is the first multi-cohort analysis of the gut microbiome in new-onset RA patients to identify consistent microbial signatures associated with drug response. This study provides valuable insights to guide future efforts in global, microbiome-informed therapy optimization.

## Results

### Study Populations

To identify microbiome features that were consistently associated with future MTX response in new-onset RA patients (NORA), we studied 3 RA cohorts. Cohort 1 (N = 47) consisted of patients consecutively enrolled from 3 rheumatology clinics at NYU and was previously studied by our group^4^. Cohort 2 (N = 24) was newly enrolled for this study at NYU and UCSF. Cohort 3 (N = 29) consisted of UK patients enrolled between April 2017 and July 2019 from 12 outpatient rheumatology clinics in England, UK; data for Cohort 3 were downloaded from Sequence Read Archive (SRA). We reasoned that the study of three distinct cohorts collected over different time spans and geographic locations would provide an opportunity to identify microbiome-based MTX response predictors that generalize across RA patients. Further, joint analysis could lead to increased sample size and power to detect associations. Demographic data for the three cohorts (N = 100) are provided in **Table 1**.

**Table 1.**
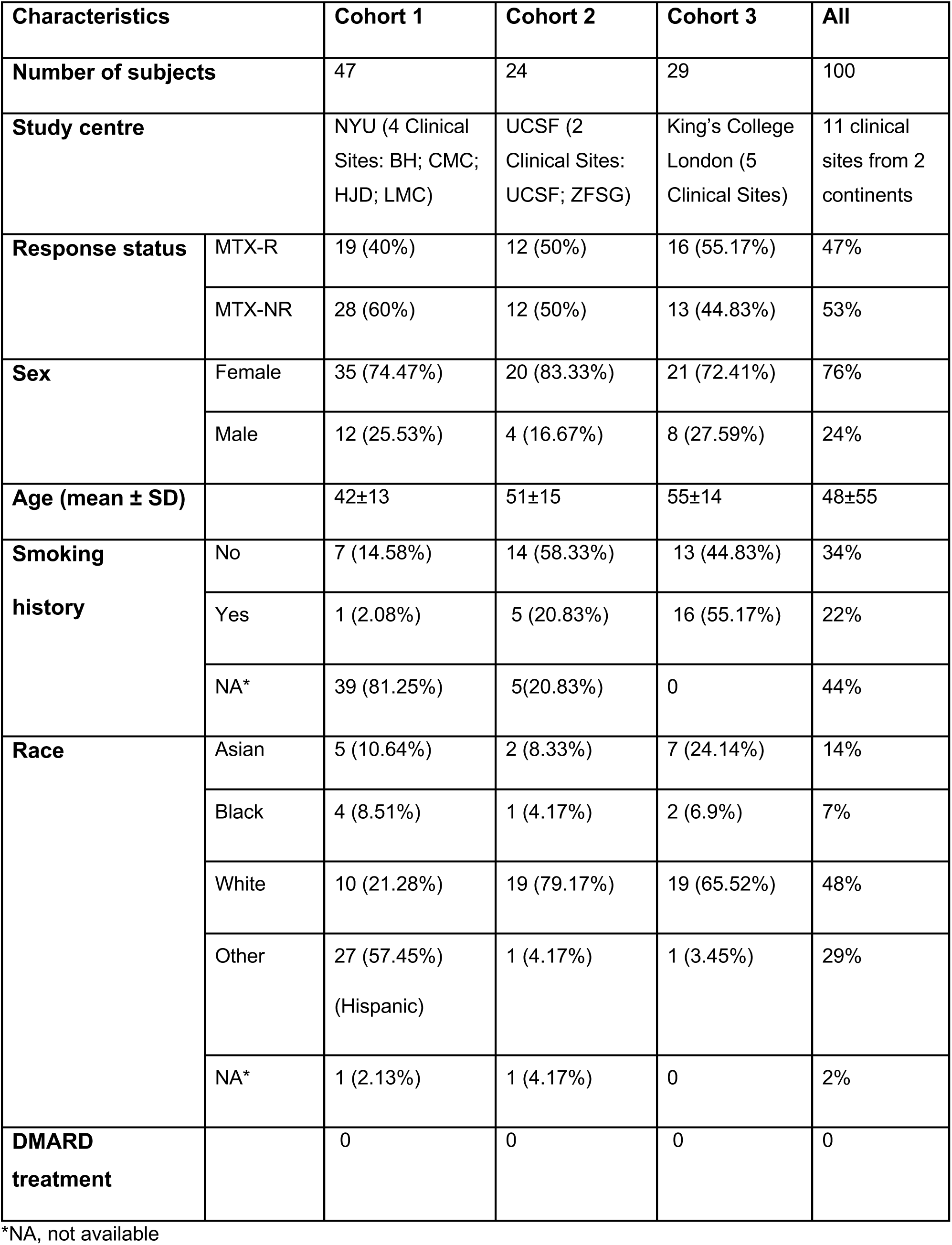
Demographic characteristics of RA patients.

All patients received oral MTX as standard-of-care treatment. MTX response was assessed 12-16 weeks after the initiation of MTX therapy. Patients were classified as MTX-R (N = 47, 47%) or MTX-NR (N = 53, 53%) as previously described^4^ based on an improvement in DAS28 ≥ 1.8 after initiation of MTX monotherapy and without the need for additional or alternative therapy.

The mean age of the study population was 48.6 ± 15.2 years, with no difference between study groups (MTX-R 48.0 ± 15.9, MTX-NR 49.0 ± 14.7, Welch’s t-test, *p =* 0.76; **Supplementary Figure 1A; Table 1; Supplementary Table 1**). Consistent with the female predominance of RA^5^, 76% of subjects were female, with similar sex distribution in MTX-NR (43 females, 10 males) and MTX-R (33 females, 14 males) (**Supplementary Figure 1B; Table 1; Supplementary Table 1**).

### Pretreatment gut microbiome is associated with future clinical response to MTX in three cohorts

We previously showed that the pre-treatment gut microbiome of NORA patients was associated with future clinical response^4^, and others have confirmed this finding^9,28^. Here we ask whether microbiome-drug response associations are cohort-specific or whether generalizable trends can be detected in a larger combined cohort of patients spanning scales of geography and time. As expected, upon joint analysis^29,30^, we found that cohort was significantly associated with microbiome structure and function (**Supplementary Figure 2**); this is potentially due to differences in sample collection and technical workflows (**Supplementary Figure 3**) as well as inherent microbiome variation associated with geographical location. To account for these cohort-specific batch effects, we used the MMUPHin tool to normalize and batch-correct the microbiome data for subsequent integrated analysis^31^. Notably, genus-level microbiome variance explained by cohort decreased markedly from 11% before correction (PERMANOVA R² = 0.109, *p* = 0.0009) to 3% after correction (PERMANOVA R² = 0.03, *p* = 0.004) (**Supplementary Figure 2**).

Using the batch-corrected dataset, we first analyzed differences in microbial taxonomic diversity. Species Shannon diversity (alpha diversity) did not differ by MTX response (Welch’s t-test, *p* = 0.06; **Supplementary Figure 4**). In contrast, community composition (beta diversity) was associated with MTX response at the phylum, genus, species and strain levels (PERMANOVA, *p* < 0.05; **Figure 1A-B**). To identify specific taxa associated with MTX response, we used multivariate linear regression, adjusting for age, sex, and cohort. At the phylum level, we found that MTX-NR exhibited a higher abundance of *Firmicutes* and a lower abundance of *Bacteroidota* (linear regression FDR ≤ 0.2; **Figure 1C; Supplementary Table 2**), similar to what we reported previously^4^. When stratified by cohort, Cohorts 1 and 3 showed concordant trends, with low-to-moderate heterogeneity (**Supplementary Figure 5A-B; Supplementary Table 2;** 𝛕^2^=0.004, H^2^=1.5, I^2^=34%, p=0.23). Next, we examined differential abundance at lower taxonomic levels, testing 79 microbial families, 159 genera, and 235 species. One family, 5 genera and 2 species were differentially abundant (linear regression, FDR < 0.2; **Figure 1D-F**; **Supplementary Table 3, 4 and 5**). The *Odoribacteriaceae* family and the *Odoribacter* genus were lower in MTX-NR (linear regression, FDR < 0.2; **Figure 1D-E**; **Supplementary Table 3 and 4**). The *Dorea* and *Ruminococcus* genera were significantly higher in MTX-NR (linear regression, FDR < 0.2; **Figure 1E**; **Supplementary Table 4**). At the species level, *Bacteroides caccae* was lower and *Ruminococcus bromii* was higher in MTX-NR (linear regression, FDR < 0.2; **Figure 1F; Supplementary Figure 5C-D; Supplementary Table 5**). In summary, these findings identify microbiome features associated with MTX response across distinct cohorts and strengthen support for the hypothesis that the gut microbiome can predict future MTX response.

**Figure 1.**
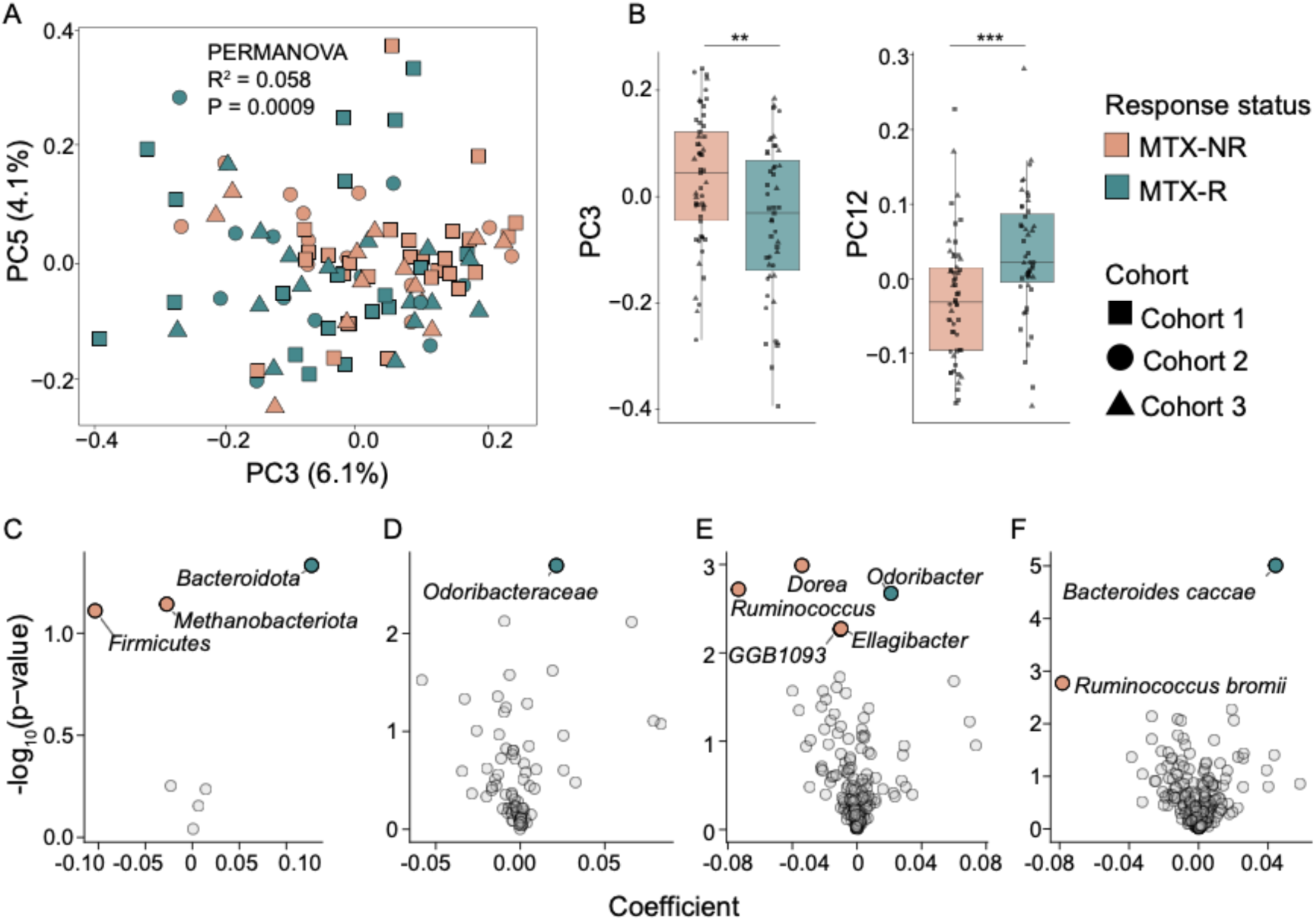
Human gut microbial taxa are associated with future MTX response across multiple pre-treatment NORA cohorts. **(A)** Principal Coordinate Analysis (PCoA) of genus-level gut microbiome profiles based on Bray-Curtis dissimilarity of proportional relative abundances from MTX-R and MTX-NR patients (PERMANOVA R² = 0.058, *p* = 0.0009). **(B)** Boxplots representing scores of PCoA component 3 and component 12. Significance is indicated as follows: \*\**p* < 0.01, \*\*\**p* < 0.001 (Wilcoxon rank sum test). Differential abundance of bacterial taxa between MTX-R and MTX-NR groups at the phylum **(C)**, family **(D)**, genus **(E)**, and species **(F)** levels. Taxonomic comparisons were performed using a linear regression meta-analysis using the MMUPHin tool, adjusting for age and gender and accounting for cohort-specific batch effects. Results are shown as a volcano plot (regression coefficient vs. significance), where colored points represent differentially abundant taxa (FDR ≤ 0.2).

### Pretreatment gut microbial function is associated with MTX response status across cohorts

Given the observed difference in microbial community composition by MTX response, we next investigated microbial functional profiles, looking at gene and pathway abundances. Whereas the total number of distinct gene functions did not differ (Wilcoxon rank-sum test *p* = 0.34; **Supplementary Figure 6**), the composition of gene functions differed by MTX response, suggesting that genes encoded by gut microbes are associated with and may influence MTX treatment outcomes (PERMANOVA, R² = 0.019, *p* = 0.031; **Figure 2A-B**). Using a multivariate linear regression adjusting for age, sex and cohort, we detected 37 genes that were differentially abundant by future response status, including 8 that were enriched and 29 that were depleted in MTX-NR (FDR ≤ 0.2; **Figure 2C; Supplementary Table 6**). These included genes involved in amino acid, carbohydrate, and nucleotide metabolism, among others, suggesting broad microbial metabolic pathways are associated with MTX response (**Supplementary Table 6**).

**Figure 2.**
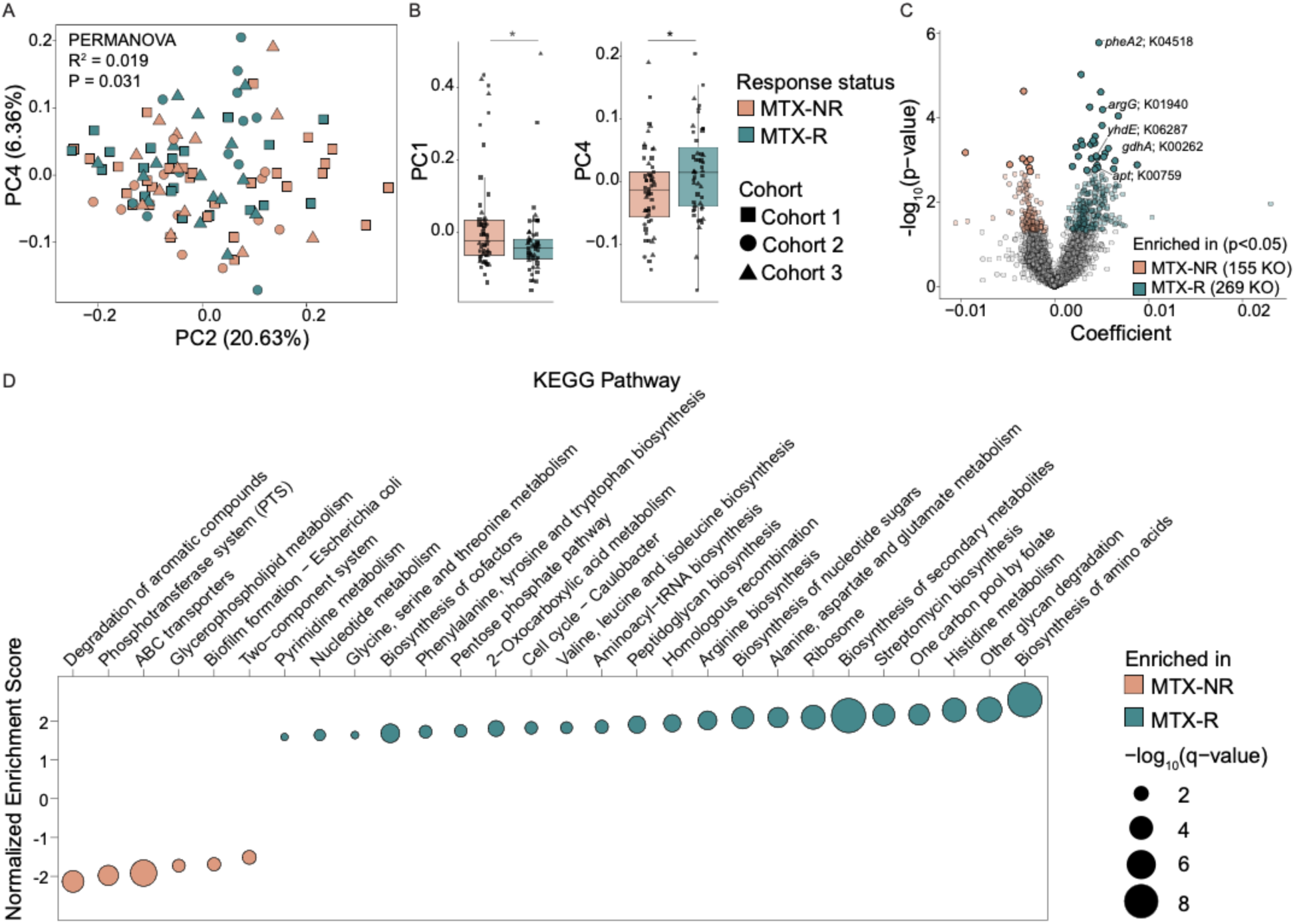
**Multiple gut microbiome functions are associated with MTX response across pre-treatment NORA cohorts**. **(A)** PCoA of gut microbiome functional profiles (KEGG ortholog abundances) based on Bray-Curtis dissimilarity in MTX-R and MTX-NR patients (PERMANOVA, R² = 0.019, *p* = 0.031). **(B)** Boxplots representing score of PCoA component 1 and component 4. Significance is indicated as follows: *p* < 0.05 (Wilcoxon rank sum test). **(C)** Differential abundance of KEGG orthologs between MTX-R and MTX-NR groups. Linear regression meta-analysis was performed with MMUPHin, adjusting for age, sex, and cohort. Results are shown as a volcano plot, where colored points represent significantly different orthologs (solid points, FDR ≤ 0.2; transparent points, *p*_unadjusted_<0.05). **(D)** Differentially enriched microbial pathways in MTX-R and MTX-NR as assessed by GSEA. The dot plot displays significantly enriched pathways, with point size representing –log₁₀(q-value) and color indicating enrichment in MTX-R or MTX-NR. Pathways with FDR ≤ 0.05 are shown.

Similarly, analysis of pathway abundances (MetaCyc pathways^32^) revealed that the pre-treatment microbiome differed by future response status. While the total number of pathways was similar between groups (Welch’s t-test, *p* = 0.22; **Supplementary Figure 7A**), we found that the composition of pathway abundances differed by MTX response (PERMANOVA, R² = 0.027, *p* = 0.018; **Supplementary Figure 7B-C**). Adjusting for age, sex and cohort, we found that 5 out of 482 pathways that were differentially abundant (FDR ≤ 0.2, **Supplementary Figure 7D; Supplementary Table 7**). Two pathways involved in purine nucleotide biosynthesis were depleted in MTX-NR: “inosine-5′-phosphate biosynthesis I” (PWY-6123) and “inosine-5′-phosphate biosynthesis II (PWY-6124) (FDR ≤ 0.2, **Supplementary Figure 7D; Supplementary Table 7**). In contrast, MTX-NR exhibited enrichment of “purine nucleobases degradation I, anaerobic” (P164-PWY), “formaldehyde assimilation III, dihydroxyacetone cycle” (P185-PWY), and “superpathway of glycerol degradation to 1,3-propanediol” (GOLPDLCAT-PWY) (FDR ≤ 0.2, **Supplementary Figure 7D; Supplementary Table 7**). Together, these findings indicate MTX-NR microbiota have reduced capacity for nucleotide synthesis and enhanced capacity for purine degradation, glycerol metabolism and formaldehyde assimilation.

In addition, we performed Gene Set Enrichment Analysis (GSEA) on ranked KEGG Ortholgs (KO) abundances to enhance sensitivity for identifying concerted shifts across biologically related gene sets^33^. A total of 29 metabolic pathways were differentially abundant (FDR < 0.05; **Figure 2D; Supplementary Table 8**), of which 7 pathways were enriched and 22 were depleted in MTX-NR. Microbiomes of MTX-NR showed depletion of metabolic pathways involved in nucleotide/cofactor metabolism, amino acid biosynthesis, protein synthesis, and cellular growth (**Figure 2D)**. Notably, several amino acid pathways depleted in MTX-NR and enriched in MTX-R —including arginine, histidine, and tryptophan metabolism—have previously been linked to arthritis and MTX efficacy^34–36^, suggesting that increased microbial amino acid metabolism may contribute to improved treatment response. Collectively, these findings suggest that the MTX-NR microbiome exhibits reduced potential for amino acid and nucleotide metabolism, central carbon processing, and the biosynthesis of cofactors and secondary metabolites, whereas these pathways are enriched in MTX-R.

### Trans-kingdom pathways associated with future MTX response

In the host, several pathways have been implicated in shaping the MTX response^37,38^, including genes involved in nucleotide metabolism, adenosine production^39^, folate-mediated reactions^40,41^, and amino acid metabolism, particularly histidine^36^, tryptophan^35^, and arginine^34^. Many of these pathways are conserved in microorganisms^7^, but whether these specific microbial pathways or genes are associated with MTX response remains unclear. Interestingly, our GSEA revealed significant enrichment of several of these pathways in MTX-R (FDR < 0.05; **Figure 2D; Supplementary Table 8**). We therefore asked whether the differentially abundant KOs identified in our dataset overlapped with genes previously associated with MTX efficacy.

We first focused on microbial functions involved in nucleotide metabolism, as this pathway is central to the mechanism-of-action of MTX^42,43^. KOs associated with adenine phosphoribosyltransferase (*apt*; K00759) and nucleoside triphosphate pyrophosphatase (*yhdE*; K06287) were significantly higher in the microbiome of MTX-R (linear regression, FDR < 0.2; **Figure 3A-B; Supplementary Figure 8A-B**). While nucleoside triphosphate pyrophosphatase (*yhdE*) broadly modulates nucleotide metabolism, *apt* specifically catalyzes the conversion of adenine to adenosine monophosphate (AMP), which can subsequently be metabolized extracellularly to adenosine^44^. Given that MTX exerts anti-inflammatory effects partly through adenosine^39,45^, enrichment of microbial nucleotide metabolism in MTX-R may enhance adenosine availability and contribute to adenosine-mediated immunoregulation.

**Figure 3.**
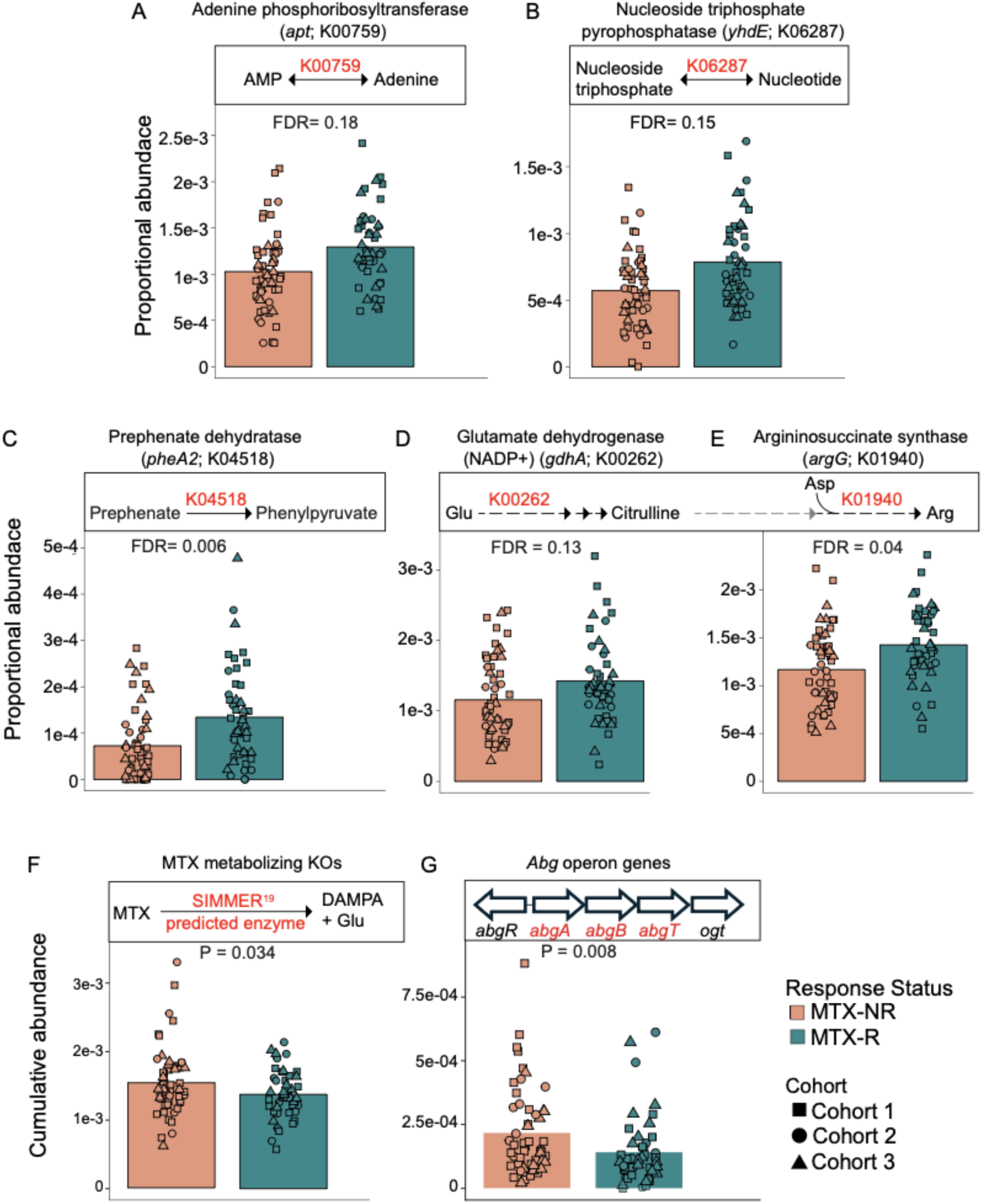
Nucleotide metabolism and amino acid synthesis pathways are depleted and MTX metabolism genes are enriched in MTX-NR. Abundance of **(A)** Adenine phosphoribosyltransferase (*apt*; K00759), **(B)** nucleotide triphosphate pyrophosphatase (*yhdE*; K06287), **(C)** prephenate dehydratase (*pheA2*; K04518), **(D)** glutamate dehydrogenase (*gdhA*; K00262), and **(E)** argininosuccinate synthase (*argG*; K01940). Statistical significance was assessed using linear regression, with FDR *p* values reported. Cumulative abundance of (**F**) 9 KEGG orthologs predicted to metabolize MTX^19^ (**Supplementary Table 9**) and of (G) *abg* operon genes (*abgA*, *abgB*, *abgT*) (**Supplementary Table 10**) across all cohorts. Data are presented cumulative abundance values. Bars represent group means, with individual samples overlaid as points. Statistical significance was assessed using a Welch’s t-test, *p* values as reported.

Given prior evidence associating amino acid metabolism to arthritis and MTX response^34,35^, we next interrogated differentially abundant KOs involved in amino acid metabolism. Consistent with prior studies^35,46^, the tryptophan biosynthesis pathway was enriched in MTX-R (**Figure 2D**). Moreover, MTX-R showed an enrichment in prephenate dehydratase (*pheA2*; K04518), which converts prephenate to phenylpyruvate in phenylalanine, tyrosine and tryptophan synthesis^47^ (linear regression, FDR = 0.006; **Figure 3C; Supplementary Figure 8C**).

Furthermore, microbial arginine metabolism differed between treatment groups. L-arginine can ameliorate arthritis, reducing disease severity and bone loss^34^. MTX-R showed an enrichment of glutamate dehydrogenase [NADP+] (*gdhA*; K00262) and arginino-succinate synthase (*argG*; K01940)(linear regression, FDR < 0.2; **Figure 3D-E; Supplementary Figure 8D-E**). Both of these enzymes are involved in L-arginine synthesis pathways ^48,49^

Altogether these observations suggest that the microbiome of MTX-R have increased capacity for nucleotide and amino acid production, potentially contributing to enhanced MTX response.

### Microbial genes implicated in MTX metabolism are associated with future MTX response

Recently, we reported that the gut microbiome modulates MTX bioavailability and therapeutic outcomes in arthritis^4,5,19^. Building on this observation, we investigated whether the microbial capacity for MTX metabolism differs between MTX-R and MTX-NR. We previously reported 18 microbial enzymes (Prokka annotations) predicted to metabolize MTX^19^; 9 of these were mapped to KO genes detected in our three cohorts (**Supplementary Table 9**). Interestingly, the cumulative abundance of these putative MTX-metabolizing KOs was significantly enriched in MTX-NR (Welch’s t-test, *p* < 0.05; **Figure 3F**), with concordant trends observed in each individual cohort (**Supplementary Figure 8F**).

Moreover, we have recently shown that gut microbes possess the ability to produce metabolites such as p-methylaminobenzoyl-L-glutamic acid (p-MABG) through the metabolism of MTX^50^. The *abg* operon mediates the transport and metabolism of a structurally related compound, p-aminobenzoyl-L-glutamic acid (p-ABG)^51^. This operon consists of three genes: *abgA* and *abgB*, which encode p-aminobenzoyl-glutamate hydrolase subunits A and B, and *abgT*, which encodes the transporter subunit T (**Figure 3G**). We therefore asked whether the KOs corresponding to genes in the *abg* operon were differentially abundant (**Supplementary Table 10**). Notably, the cumulative abundance of KOs corresponding to these three genes was significantly higher in MTX-NR (Welch’s t-test, *p* = 0.008; **Figure 3G**), with similar effects observed across the cohorts (**Supplementary Figure 8G**).

Altogether, these findings suggest that the MTX-NR microbiome is enriched with genes involved in MTX degradation. Microbial MTX-degrading genes may reduce MTX bioavailability, thereby contributing to poor response to MTX.

### Baseline DAS28 score is associated with future clinical response in three cohorts

Finding that the pretreatment microbiome of RA patients was associated with MTX response, we next asked if a predictive model could be developed using microbiome features, with or without the inclusion of clinical parameters. First, we asked if clinical parameters, such as age, sex, baseline disease activity, or serologic status were associated with future MTX response, as prior studies have suggested some associations^52–55^. In patients with established RA, baseline DAS28-ESR, anti-citrullinated protein antibody (ACPA), and Health Assessment Questionnaire (HAQ) scores have been identified as top predictors of favorable MTX response^56^. Whether this is also the case in new-onset rheumatoid arthritis (NORA) remains unknown.

Interestingly, in NORA patients, we observed that baseline DAS28 scores were higher in future MTX-R compared with MTX-NR (Welch’s t-test, *p* = 0.012; **Figure 4A; Supplementary Table 1**). This trend was observed across all three cohorts, suggesting that higher disease activity at baseline is associated with an increased likelihood of patients responding to MTX. We next asked if baseline DAS28 was correlated with pre-treatment microbial abundances. This would enable us to identify taxa that are linked to disease activity and may impact treatment outcomes. At the genus level, 4 bacterial genera were significantly correlated with baseline DAS28 (Spearman’s rank correlation test, FDR ≤ 0.2; **Figure 4B; Supplementary Figure 9A; Supplementary Table 11**), with 3 showing positive correlations (*Allisonella, Ligilactobacillus,* and *Segatella)*. At the species level, we found *Eubacteriaceae bacterium* was negatively associated with baseline DAS28 (Spearman’s rank correlation test, FDR = 0.13; **Supplementary Figure 9A; Supplementary Table 11**). In contrast, 9 bacterial species were positively correlated with DAS28, including *Allisonella histaminiformans, Clostridium fessum, Dialister hominis, Gemmiger SGB15295, Lachnospiraceae bacterium OF27 8pH9A, Ligilactobacillus ruminis, Segatella brunsvicensis, Segatella copri,* and *Veillonella atypica* (Spearman’s rank correlation test, FDR ≤ 0.2; **Figure 4C; Supplementary Figure 9B; Supplementary Table 12**). Notably, *Segatella copri,* formerly classified as *Prevotella copri,* has been repeatedly linked to RA pathogenesis^52–55,57^. Our findings extend these observations by further associating *S. copri* abundance with disease severity.

**Figure 4.**
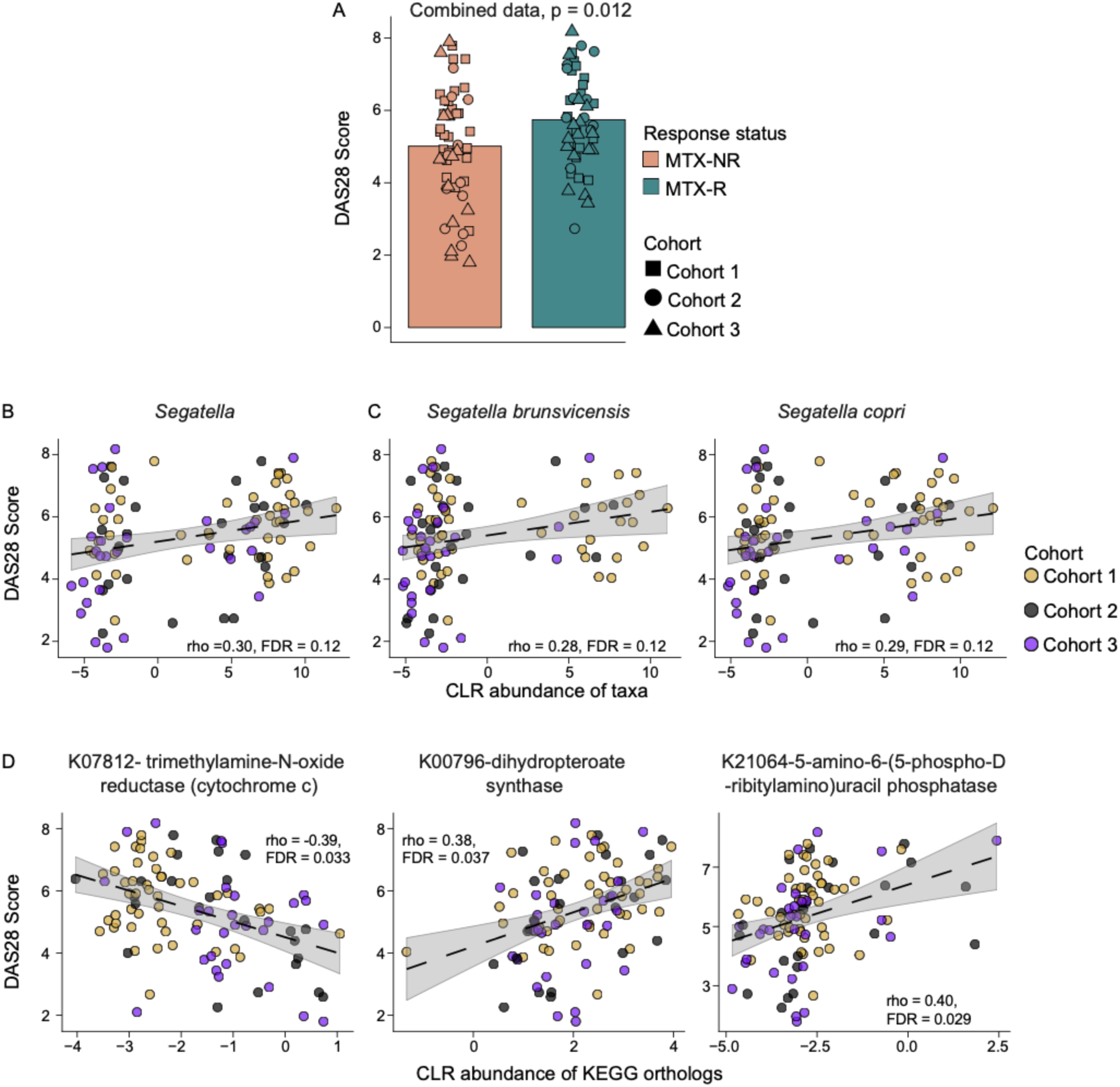
Baseline DAS28 scores are associated with bacterial genera, species and gene families in NORA subjects. **(A)** MTX-NR exhibits lower DAS28 scores at baseline compared to MTX-R in the combined analysis. Bars represent group means, with individual samples overlaid as points. Statistical comparisons were performed using a Welch’s t-test, with *p*-values indicated. Spearman’s rank correlation was performed between DAS28 scores and the CLR abundance of **(B)** bacterial genera, **(C)** species and (**D**) KEGG orthologs. The y-axis represents DAS28 scores and x-axis represent microbial taxa or KEGG features.

We next investigated whether microbial functions are associated with DAS28 scores. Spearman correlation analysis revealed 495 KOs were significantly correlated with DAS28, including 461 positive and 34 negative correlations (FDR ≤ 0.2; **Figure 4D; Supplementary Figure 9C; Supplementary Table 13**). Notably, trimethylamine-N-oxide (TMAO) reductase [cytochrome c] (*torZ*; K07812) was negatively associated with DAS28 (FDR = 0.033; **Figure 4D**). Given that TMAO is generally pro-inflammatory^58,59^, this suggests that bacterial TMAO-reducing activity may lower systemic TMAO levels, potentially contributing to lowering disease severity in RA patients. Among the positively correlated KOs, 2 KOs were linked to distinct vitamin biosynthesis pathways (Spearman’s rank correlation test, FDR ≤ 0.2; **Figure 4D; Supplementary Table 13**): dihydropteroate synthase (*folP*; K00796) has been shown to catalyze a key step in the *de novo* folate pathway^60^ and 5-amino-6-(5-phospho-D-ribitylamino)uracil phosphatase (*ycsE/yitU/ywtE*; K21064) functions in riboflavin biosynthesis^61^. This suggests that microbial capacity to produce folate and riboflavin is associated with elevated disease severity at baseline.

### Pretreatment gut microbiome predicts MTX response across cohorts

Given the aforementioned associations, we next asked whether these baseline clinical and microbial features could be used to develop a cross-cohort predictive model using random forests. We trained and validated random forest models incorporating age, sex, DAS28, and/or KEGG abundance features for predicting MTX response, with performance evaluated by the area under the receiver operating curve (AUC-ROC).

We first performed a leave-one-cohort-out (LOCO) framework (see Methods) to evaluate generalizability on an independent validation cohort: models were trained on two cohorts and tested on the remaining third cohort. Model performance using KEGG orthologs was dependent on training cohort size and geographic representation: a model developed using Cohorts 1 and 3 (US and UK combined N = 76) performed with an AUC of 0.8 on Cohort 2 (US cohort N = 24), whereas model performance was lowest for Cohort 3 (AUC=0.59), in which the model was trained on US patients (N=71) and validated on UK patients (N=29) (**Figure 5A**). Overall, when averaging performance across 3 independent validation cohorts, KEGG-based model performance was moderate, achieving an AUC of 0.69 (**Figure 5A**). The model based on KEGG features (**Supplementary Table 14**) was more predictive than those based on clinical parameters (age, DAS28 score, sex) (Mean AUC=0.56; **Figure 5A**). Inclusion of clinical parameters alongside KEGG features did not improve model performance (Mean AUC = 0.68; **Figure 5A**). These findings suggest that the human gut microbiome provides cross-cohort predictive power and can exceed performance achieved by models built using clinical parameters alone.

**Figure 5.**
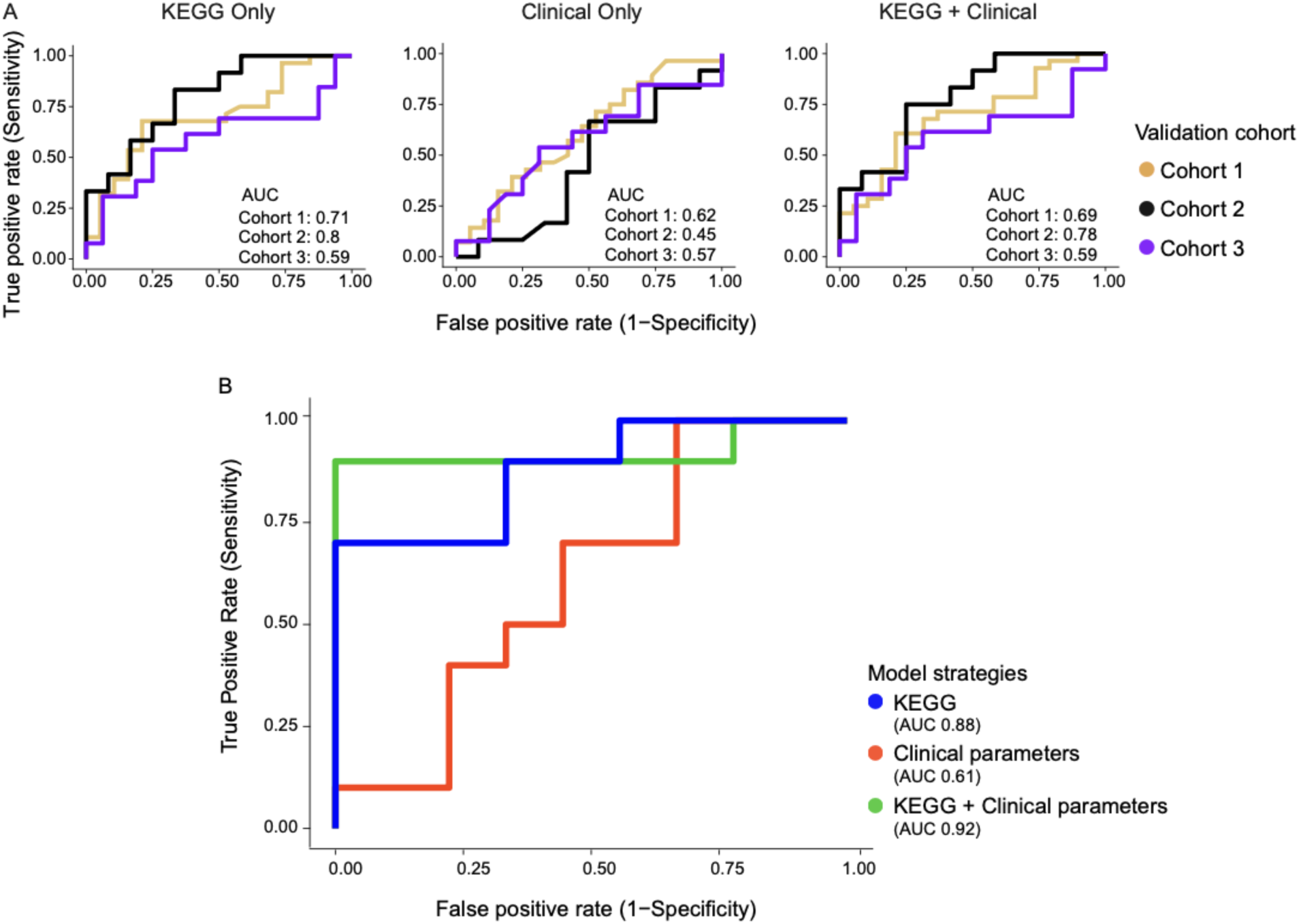
Microbiome based model outperforms clinical parameter based model in predicting MTX response in patients. Random forest model was built using Boruta selected KEGG orthologs alone, clinical parameters (DAS28, age and sex) or their combinations (KEGG + Clinical) under two different frameworks. Models were either trained on (**A**) two cohorts and validated on a held-out third cohort (LOCO framework), or (**B**) using ∼80% of samples for training and the remaining ∼20% were for validating while balancing the response groups (stratified split framework). The area under the curve (AUC) for each model is shown. Models shown correspond to iterations in which the KEGG-only strategy achieved the best performance across 50 iterations. Lines represent performance on individual test cohorts in the LOCO framework and KEGG-only, clinical-only, or combined models in the stratified split framework.

Given that training cohort size and geographic representation was linked to model performance in LOCO analysis, we next asked whether a combined training cohort, inclusive of patients from all three cohorts, would have greater predictive power. We applied a stratified train-validation framework (see Methods; **Supplementary Table 15**) in which we pooled all three cohorts and then split them into 80/20 training and validation cohorts: patient samples were divided into a training cohort (N=81; 38 MTX-R and 43 MTX-NR) and a held-out validation cohort (N = 19; 9 MTX-R and 10 MTX-NR; **Supplementary Table 15**). We applied the Boruta algorithm^62^ on the training cohort for feature selection, resulting in identification of 13 KOs that were predictive of MTX response (**Supplementary Table 16**). During cross-validation on the training samples, models incorporating KEGG features demonstrated strong discriminatory performance (**Supplementary Table 17**): the KEGG-only model achieved a mean AUC of 0.879 ± 0.078, while the combined KEGG features with clinical variables showed the highest performance (AUC = 0.895 ± 0.071). In contrast, clinical-only models exhibited moderate performance (AUC = 0.63), and individual variables such as DAS28, age, or gender performed close to random classification (AUC < 0.59).

When tested on the held-out validation cohort, random forest modeling using KEGG features alone achieved similarly high predictive performance (AUC = 0.88; **Figure 5B; Supplementary Table 17**), and outperformed models based on clinical variables such as DAS28, age, or gender (AUC ≤ 0.67; **Figure 5B; Supplementary Figure 10; Supplementary Table 17**).

Adding clinical features individually to Boruta-selected KEGG features (KEGG + Age, KEGG +Sex or KEGG + DAS28) slightly reduced predictive performance (AUC = 0.84-0.87) (**Supplementary Figure 10**), and a model combining all clinical and KEGG features showed modest improvement in performance over KEGG-only features (AUC = 0.92) (**Figure 5B**). Altogether, microbiome based models achieved robust discriminative performance in predicting MTX response across cohorts.

To better understand the microbiome features that were predictive of future MTX response, we next examined the 13 KOs that were selected by Boruta and found that these KOs represented diverse microbial functions with plausible links to RA and/or autoimmunity. First, K01197 (*hya*; hyaluronoglucosaminidase) encodes an enzyme that breaks down hyaluronic acid (HA), a key component of the gut extracellular matrix^63,64^ as well as the joint synovium^65,66^. HA serves as a microbial substrate that supports the growth of specific taxa^67^. Importantly, reduced synovial HA levels have been linked to RA^68^.

Second, 5 ribosomal proteins were also predictive: 3 proteins of the large subunit (K02892, *rplW*, large subunit ribosomal protein L23; K02874, *rplN*, large subunit ribosomal protein L14; K02907, *rpmD*, large subunit ribosomal protein L30) and 2 of the small subunit (K02952, rpsM, small subunit ribosomal protein S13; K02965, *rpsS*, small subunit ribosomal protein S19). Notably, K02892 (*rplW*), is linked to RA pathology^69^: its eukaryotic homolog RPL23A, is a ubiquitously expressed autoantigen that elicits both T cell and autoantibody responses specifically in RA patients^69^. Additionally, the eukaryotic homolog of K02874 (*rplN*) has been suggested as an autoantigen in autoimmune disease^70^. Further, these were enriched in future MTX-R (**Supplementary Table 6**) and/or positively correlated with DAS28 (**Supplementary Table 13**). The remaining 7 KOs lacked clear links to RA and were related to aminoacyl-tRNA biosynthesis (K00773, K01876), nutrient acquisition (K01995, K04758), protein degradation (K01358), RNA turnover (K08300), and DNA repair (K09760) (**Supplementary Table 16)**. Overall, our study suggests that a subset of KOs implicated in autoimmune disease pathogenesis may also serve as predictors of treatment response.

Taken together, these results show that the pretreatment microbiome can predict future MTX response in an independent validation cohort, as demonstrated by LOCO analysis. Furthermore, a model that is inclusive of the diversity of RA patients affords greater predictive power.

## Discussion

Multiple studies have demonstrated links between the human gut microbiome and treatment outcomes^4,9,16–18^, but whether predictive microbial features are concordantly predictive across different study populations is unclear^17,18^. Our study represents the first integrated analysis of the gut microbiome in NORA patients to identify cross-cohort microbial signatures associated with MTX response and provides a framework for developing a generalizable model to predict treatment response. By jointly analyzing 3 cohorts from the US and UK, we identified microbial taxa and functions that are associated with and predict MTX response across cohorts, and many of these show biologically meaningful links to RA or MTX.

Consistent with our previous findings^4^, *Bacteroidota* were depleted and *Firmicutes* were enriched MTX-NR in the combined cohort. This enrichment of *Firmicutes* is notable because we previously reported that this phylum is enriched for MTX metabolizing species: 62% (10 out of 16 tested) of *Firmicutes* depleted MTX from the culture medium compared to 0% (0 out of 14) of *Bacteroidota*^19^. In contrast, members of the *Bacteroidota* phylum tend to be sensitive to MTX treatment as assessed by growth inhibition^7^. Together, these observations lead us to speculate that higher abundance of *Firmicutes* in MTX-NR may lead to increased MTX metabolism, thereby leading to reduced likelihood of adequate response. In contrast, higher levels of sensitive *Bacteroidota* in MTX-R may allow MTX to exert its anti-inflammatory effects by acting on both the microbiota and the host to reduce inflammation.

In furtherance of this possibility, the combined analysis demonstrated that candidate MTX metabolizing genes are enriched in MTX-NR, with concordant trends observed in all three cohorts. These candidate MTX metabolizing genes were identified using a similarity algorithm, SIMMER^19^, and in our previous work, we showed that these genes were higher in MTX-NR in Cohort 1^19^. This result is now reproduced here in two additional patient cohorts. Furthermore, these genes were enriched in mice showing increased microbiota-dependent MTX metabolism in pharmacokinetic studies^5^, and they were associated with MTX response in an SKG mouse model as assessed by clinical arthritis scores^5^. Combined with our *ex vivo* studies of RA patient microbial communities showing that MTX depletion from the media is inversely correlated with future MTX response^4^, these findings provide strong, quantitative orthogonal lines of evidence that microbial metabolism of MTX by the human gut microbiome is clinically relevant. Indeed, it may be a modifiable factor that can be leveraged to both predict and improve treatment outcomes in RA.

Functional analysis revealed that pathways previously linked to RA and MTX in the host were also detected in the gut microbiome and associated with MTX response. Experimental evidence suggests that host metabolism of histidine^36^, tryptophan^35,46^, arginine^34^, and nucleotides^39,44^ are mechanistically relevant to MTX response. In our analysis, all of these gut microbial pathways were differentially abundant between MTX-R and MTX-NR. These findings provide evidence linking microbial correlates of host metabolic processes to MTX response in RA. Experimental studies to assess whether these microbial pathways contribute to MTX response are needed.

We unexpectedly found that baseline DAS28 scores were significantly higher in future MTX-responder subjects, a trend that was reproducible across all three cohorts, suggesting higher disease activity at baseline is associated with increased likelihood of favorable MTX response in new-onset RA patients; this is in contrast to findings in patients with established RA^56^. Nevertheless, predictive models using DAS28 achieved only moderate discrimination (AUC=0.64). Interestingly, *S. copri* was positively correlated with DAS28, consistent with prior studies linking *S. copri* expansion to increased risk and severity of RA^52–55,57^. Interestingly, *Segatella*-dominated healthy omnivore individuals have been reported to exhibit elevated basal levels of serum trimethylamine *N*-oxide (TMAO), a metabolite associated with pro-inflammatory effects and atherogenesis^71^. Notably, we observed a negative correlation between DAS28 and microbial TMAO reductase (*torZ*; trimethylamine-N-oxide reductase [cytochrome c]), suggesting potential mechanistic link between TMAO reducing bacteria and lower disease activity in RA patients.

Finally, a machine learning model trained on pretreatment KEGG ortholog abundances was able to accurately predict future MTX response. Both the leave-one-cohort-out (LOCO) and stratified split frameworks revealed that microbiome-feature-based models could outperform clinical features in predicting MTX response. LOCO analysis revealed that a microbiome-based model can predict response in an independent validation cohort with high predictive power: Cohort 2, a new cohort enrolled for this study, was predicted with AUC=0.80. While the other cohorts were not similarly predicted with high power, this may be due to reduced training cohort size, class imbalance in response status^72,73^, and/or training cohorts that were not representative of the validation cohort^74^, suggesting that training on larger, more representative cohorts may improve cross-cohort generalizability. Accordingly, when we trained a model using samples from all three cohorts, the model performance improved on the held-out validation cohort.

Remarkably, our study identified specific KEGG orthologs linked to autoimmune disease and host-microbiota interactions. Notably, *hya* (K01197) encodes a hyaluronoglucosaminidase that acts on host hyaluronic acid (HA), a major component of the gut extracellular matrix. By degrading HA, this enzyme facilitates bacterial access and adhesion to the mucosal layer^63,64,66^, thereby influencing the epithelial barrier functions^67^ and immune signaling^75^. Interestingly, this gene appeared in two of three LOCO models (**Supplementary Table 14**) and in the stratified split model (**Supplementary Table 16**). Similarly, *rplN* (K02874) was detected in two LOCO models (**Supplementary Table 14**) and in the stratified split model (**Supplementary Table 16**). Notably, eukaryotic homologs of *rplN* (K02874) and another predictor *rplW* (K02892) are known autoantigens in patients and mouse models of autoimmune disease^69,70^. Their enrichment in MTX-R and/or positive correlation with baseline DAS28 tempts us to speculate that gut microbial proteins mimicking host autoantigens may drive inflammation in some RA patients; their enrichment serves as favorable predictors of MTX treatment response in RA, given that MTX may act on the gut microbiome to alleviate inflammation^7^.

Nevertheless, this study has limitations. First, the overall sample size was modest (N = 100), but given the challenges inherent to prospectively enrolling new onset RA patients before treatment initiation, this represents the largest cohort of RA patients to date interrogating microbiome predictors of MTX response. Second, the current analysis is primarily associative in nature, but our previous studies in mice and *in vitro* and *ex vivo* support the overall findings^4,5,19^.

Third, our final model was trained on subjects from all three cohorts, and an independent validation cohort would further strengthen our findings.

Despite these limitations, our multi-cohort analysis provides strong support for the hypothesis that the gut microbiome of RA patients is associated with and predicts future response, and that a generalizable microbiome-based predictor may advance precision medicine in RA treatment. Moreover, our study provides evidence from multiple clinical cohorts suggesting that enrichment of MTX metabolizing genes in the gut microbiome is associated with reduced likelihood of MTX response. These findings provide a foundation for future mechanistic investigations to elucidate the role of microbial metabolic pathways in modulating response to MTX.

## Methods

### Patients

Metagenomes of a total of 100 treatment-naïve RA subjects were analyzed in the study. Participants were recruited from New York University Langone Medical Center, Lutheran Hospital (Staten Island), and the University of California, San Francisco. Patients enrolled between January 2018 and December 2021 were designated as Cohort 1 (N = 47) and reported in our prior publication^4^. Patients that were not previously studied and were recruited as a part of this study between January 2018 and December 2023 formed Cohort 2 (N = 24). We also included a publicly available metagenomic dataset^9^ of treatment-naïve, newly diagnosed RA patients recruited between April 2017 and July 2019 (NCBI SRA PRJNA957107). This cohort is referred to as Cohort 3 (N = 29) and consisted of UK patients enrolled between April 2017 and July 2019 from 12 outpatient rheumatology clinics in England, UK. MTX response was defined as an improvement of ≥ 1.8 in DAS28 at 12-16 weeks post-MTX monotherapy. Clinical and demographic characteristics of each cohort are provided in **Table 1** and **Supplementary Table 1**.

### DNA library construction and sequencing

For Cohort 2, DNA was extracted using ZymoBiomics 96 MagBead DNA Kit (Cat. No. D4308) with 2 cycles of 5 minutes bead-beating. 10-80 ng of DNA was used in the Nextera DNA Flex library prep kit along with index barcodes to assemble the metagenomic library. Quality and quantity checks were performed with Quant-iT Picogreen dsDNA Kit (Invitrogen Cat. No. P7589) with equimolar pooling of samples passing quality checks. The pooled library was quality-checked using TapeStation 4200 (Agilent), followed by sequencing using an S1 flow cell on NovaSeq 6000 system (Illumina).

### Read pre-processing and profiling

All three RA metagenomic datasets were processed from raw reads using a uniform pipeline and up-to-date metagenomic analysis pipeline, as outlined below (**Supplementary Figure 3**). Demultiplexed FASTQ files were processed for adapter trimming and quality filtering with Fastp^76^. On average, 97.17 ± 2.2% of reads passed quality filtering, yielding samples with average 35.11 ± 23.16 million high quality reads per sample (**Supplementary Table 18**).

Host reads were removed by mapping to the human genome (GRCh38) with BMTagger (ftp://ftp.ncbi.nlm.nih.gov/pub/agarwala/bmtagger/). Taxonomy was annotated with MetaPhlAn (version 4.2.2) ^77^ using the CHOCOPhlAnSGB (version 202503) database.

For functional profiling, the same set of paired-end FASTQ files were merged into single files for each sample. These were then processed using HUMAnN (version 4.0 alpha.1) to determine the relative abundance of microbial pathways in different gut metagenomes ^78^. UniRef90 gene family abundances output from HUMAnN were further mapped to the Kyoto Encyclopedia of Genes and Genomes (KEGG) Orthologous groups (KOs). The fastq files are available on the NCBI Sequence Read Archive (BioProject SUB16070656).

### Quality Control

Prior to quality control, the taxonomic dataset included 21 microbial phyla, 407 families, 1,256 genera, 2028 species. Taxa were retained if their relative abundance was exceeding > 0.005% in at least 10% of samples in at least 1 cohort. After filtering, the final taxonomic profile used for all downstream analyses consisted of 7 bacterial phyla, 79 families, 159 genera, and 235 species. The same prevalence and abundance criteria were applied to KO features. Of the initial 6,518 KOs, 3,900 passed the filtering thresholds and were included in subsequent functional analyses.

### Batch correction

All statistical analysis was performed in R (version **4.4.3**). Taxonomic and functional profiles were normalized by total sum scaling to obtain relative abundances and further batch corrected with MMUPHin (version 1.20.0)^31^ to reduce cohort-specific effects that may arise from technical and biological variation and to facilitate meta-analysis. MMUPHin is a batch correction method that employs an empirical Bayes approach to model read counts with respect to batch variables and biologically relevant covariates such as age and sex of the subjects. The batch corrected data-matrix used for all the downstream analyses

### Statistical analysis of taxonomic profiles

Taxonomic diversity analysis was performed using the phyloseq package (version 1.50.4)^79^. Shannon diversity index was calculated with the *estimate_richness* function. Beta diversity was assessed using Bray-Curtis dissimilarities computed from the batch-corrected matrix with the *distance* function. The resulting matrix was used to perform principal coordinate analysis (PCoA) to visualize differences in microbial community composition among samples. Welch’s t-test was applied to the boxplots to determine whether the groups differed significantly along individual ordinations.

To evaluate differences in community composition, Permutational Multivariate Analysis of Variance (PERMANOVA) was performed on Bray-Curtis distances using the *adonis2* function from the vegan package (version 2.7.2)^33^. Boxplots on the PCoA axes represent PCoA component values. For all boxplots, top and bottom hinges correspond to the first and third quartiles, respectively, horizontal lines denote the median, and whiskers extend to the maximum and minimum values.

To identify microbial taxa that were differentially abundant between responder groups, a meta-analysis framework was applied using the MMUPHin (version 1.20.0) package. Differential abundance was assessed using feature-wise linear models, with the *lm_meta* function under the model specification: *abundance ∼Response_status +age + gender,* with cohort included as a batch variable, this model considers MTX response as the primary predictor while accounting for age and gender as covariates. Taxa showing statistically significant associations (Benjamini-Hochberg FDR ≤ 0.2) were visualized in a bar plot.

### Statistical analysis with functional profile of microbiome

Differences in KO composition were assessed using Bray-Curtis dissimilarities of relative abundances from the batch-corrected matrix and marginal boxplots were added to illustrate the distribution of sample scores along each axis. To evaluate differences in community composition, PERMANOVA was employed.

To identify KOs differentially abundant between responder groups, a meta-analysis framework was applied using the MMUPHin R package. Differential abundance was assessed using feature-wise linear models of proportionally normalized relative abundances while accounting for age and gender as covariates. KOs with FDR < 0.2 are reported.

### Gene set enrichment analysis

A rank metric for each KO was calculated as the signed significance, defined as the sign of the regression coefficient multiplied by the negative log_10_ of the p-value^80^, based on output from the MMUPHin meta-analysis. KOs were ranked in descending order to generate a pre-ranked list for gene set enrichment analysis (GSEA) using the clusterProfiler (version 4.14.6) R package^81^.

Normalized enrichment scores (NES) were computed and pathways with FDR ≤ 0.2 were considered significant. Human disease related pathways (e.g. COVID19) are excluded from figures, given our focus on the microbiome, but included in supplementary tables.

### Correlation analysis

Spearman’s rank correlation test was applied to evaluate the correlation between DAS28 score and centered log ratio (CLR)-transformed relative abundance of microbial genera and species. *p*-values were adjusted for multiple testing using Benjamini-Hochberg (BH) false discovery rate (FDR) correction, and taxa with FDR ≤ 0.2 were considered significant. Scatter plots with linear regression fits were generated for each significant taxon. For gene function correlations, CLR-transformedKO abundance was correlated with DAS28 score using Spearman’s rank correlation test with BH-adjusted FDR ≤ 0.2 considered significant.

### Identification of a microbiome functional features to predict response to MTX

We sought to build a model using the baseline KO abundance to predict MTX response. Two complementary frameworks were employed. First, a leave-one-cohort-out (LOCO) framework was employed to evaluate cross-cohort generalizability. In this strategy, a random forest model was trained on two cohorts and performance was evaluated on the held-out third cohort. Second, a stratified split framework was employed on the combined dataset. Samples were stratified by response status while maintaining cohort-specific representation (training cohort, N=81, 38 MTX-R and 43 MTX-NR; validation cohort: N=19; 9 MTX-R and 10 MTX-NR; **Supplementary Table 15**).

In both frameworks, KEGG features were selected by applying the Boruta algorithm from boruta R package (version 9.0.0) on the training cohort(s)^62^ with a maximum of 100 iterations. Tentative features were either rejected and or retained based on median importance using the *TentativeRoughFix* function. Confirmed features were used for downstream analyses. Random forest models (500 trees, 5-fold cross-validation) were then trained on selected features and evaluated on the validation cohorts. Model performance was evaluated using area under the receiver operating curve (ROC). In both frameworks, to account for variability arising from random train-validation splits and/or other intrinsically random elements of machine learning (e.g. feature selection by Boruta tool), the final model was selected as the best performing model for KEGG only strategy across 50 iterations as previously described^17,82^, in order to facilitate biological interpretation of predictive microbiome features. Performance was assessed using mean AUC (across three cohorts) for the LOCO framework and AUC for the combined cohort in microbiome-based models.

## Author Contributions

Conceptualization: RLB, JUS, PJT, RRN. Funding acquisition: JUS, PJT, RRN. Investigation: RLB, RBB, KT, DAO, JUS, PJT, RRN. Methodology: RLB, KT, PJT, RRN. Project administration: RRN. Resources: RBB, JUS, PJT, RRN. Supervision: PJT, RRN. Validation: RLB, RRN. Visualization: RLB, RRN. Writing – original draft: RLB, RRN. Writing – review & editing: all authors.

## Acknowledgements

We would like to thank members of the Nayak Lab for their helpful comments and suggestions on the manuscript. We would like to thank the patients for their contribution to science and the advancement of medicine. This work was supported by the following: R.B.B. was supported by NIH/NCATS UL1TR001445. This was work was supported by multiple grants: R01CA255116 and R01HL122593 (P.J.T), R01AR074500 (J.U.S and P.J.T), UCSF Breakthrough Program for Rheumatoid Arthritis-related Research (BPRAR; partially funded by the Sandler Foundation; P.J.T), Arthritis Foundation Center of Excellence (R.R.N); K08AR073930 (R.R.N); R03AR082036 (R.R.N.), I01CX002557 (R.R.N.), R35GM151349 (R.R.N.), Russell/Engleman Rheumatology Research Center (R.R.N.), and the Arthritis National Research Foundation (R.R.N.), Benioff Center for Microbiome Medicine (R.R.N. and P.J.T). P.J.T is a Biohub, San Francisco, Investigator.

## Supplementary Figures

**Supplementary Figure 1.**
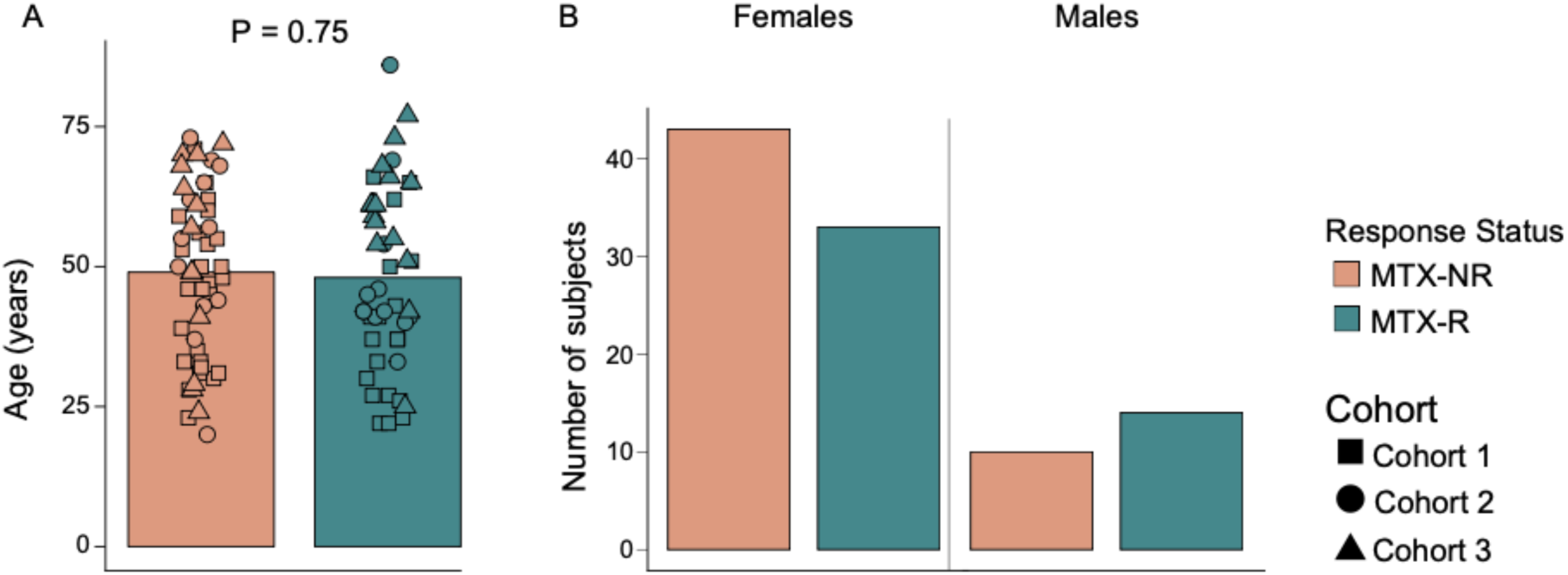
Distribution of age and sex across study cohorts. **(A)** Age did not differ between treatment groups. Data are shown as bar plots, with bars representing group means and individual samples overlaid as points. Statistical comparisons were performed using Welch’s t-test, and *p*-value reported. **(B)** Sex distribution across treatment groups shown as bar plots representing the number of males and females.

**Supplementary Figure 2.**
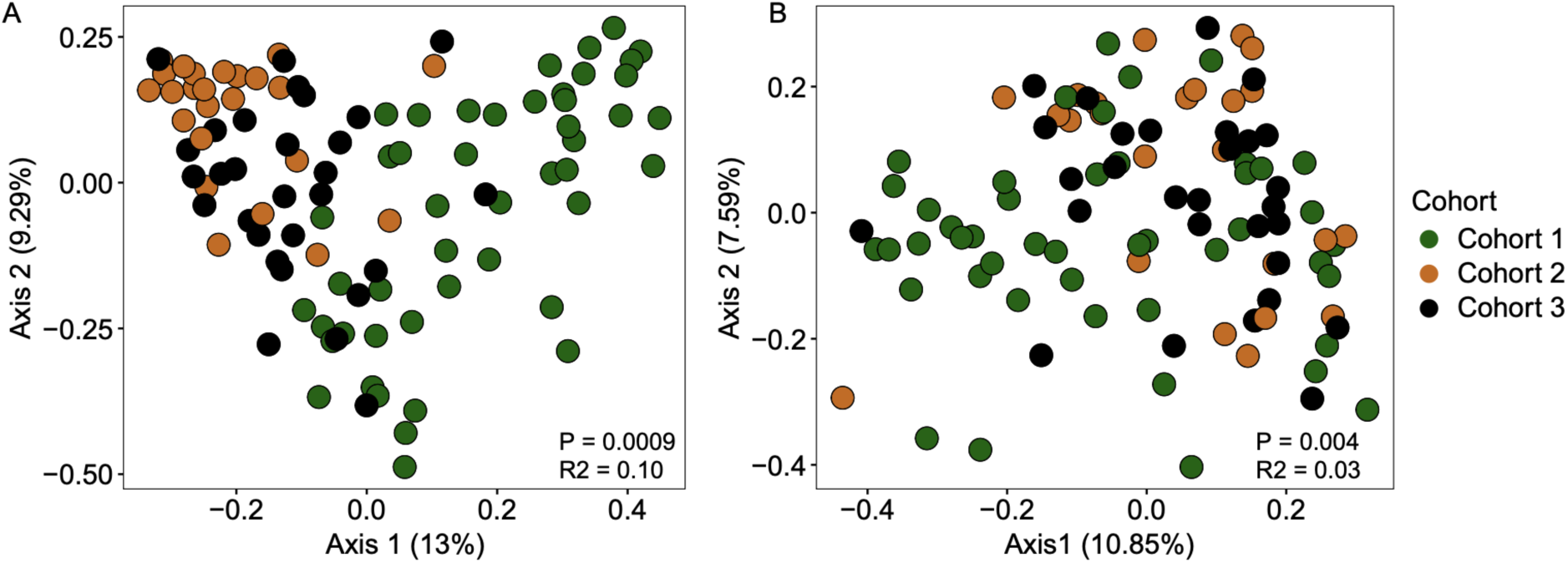
MMUPHin-normalized microbiome differences between cohorts. (**A**) PCoA of gut microbiome composition based on Bray-Curtis dissimilarity of proportional relative abundances across cohorts before MMUPHin normalization (PERMANOVA, R² = 0.10, *p* = 0.0009). (**B**) PCoA after normalization to account for cohort effects on the microbiome (PERMANOVA, R² = 0.004, *p* = 0.03).

**Supplementary Figure 3.**
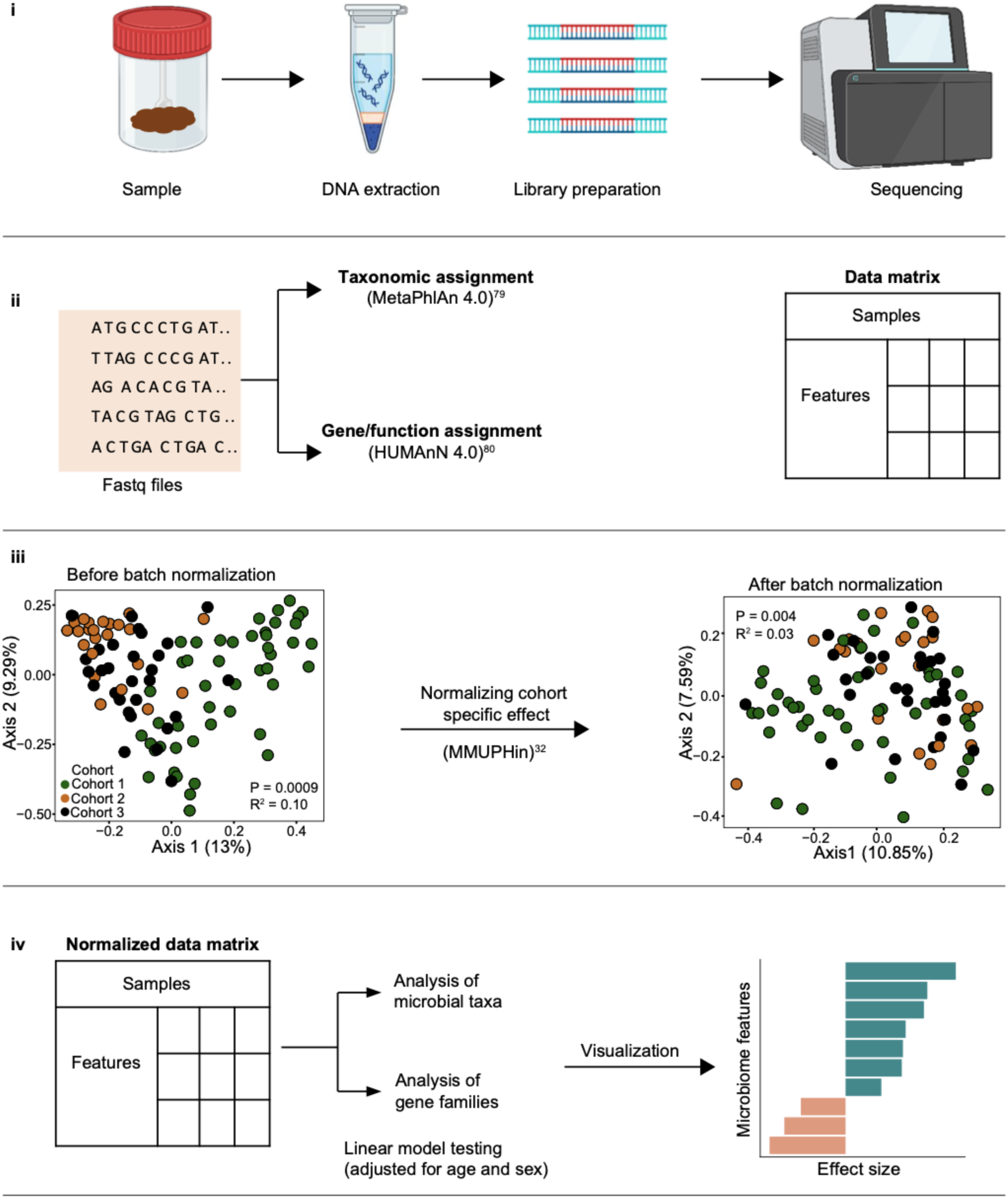
Metagenomic analysis workflow. (**i**) Shotgun metagenomic sequencing was performed on pretreatment gut microbiome samples. Each cohort underwent different DNA extraction and library preparation workflows, which is typical in microbiome studies given that there is no standard protocol. (**ii**) Taxonomic and functional profiling was conducted using MetaPhlAn and HUMAnN. (**iii**) Data were harmonized to correct for cohort-specific effects. (**iv**) Harmonized data were analyzed using linear models adjusted for age and sex.

**Supplementary Figure 4.**
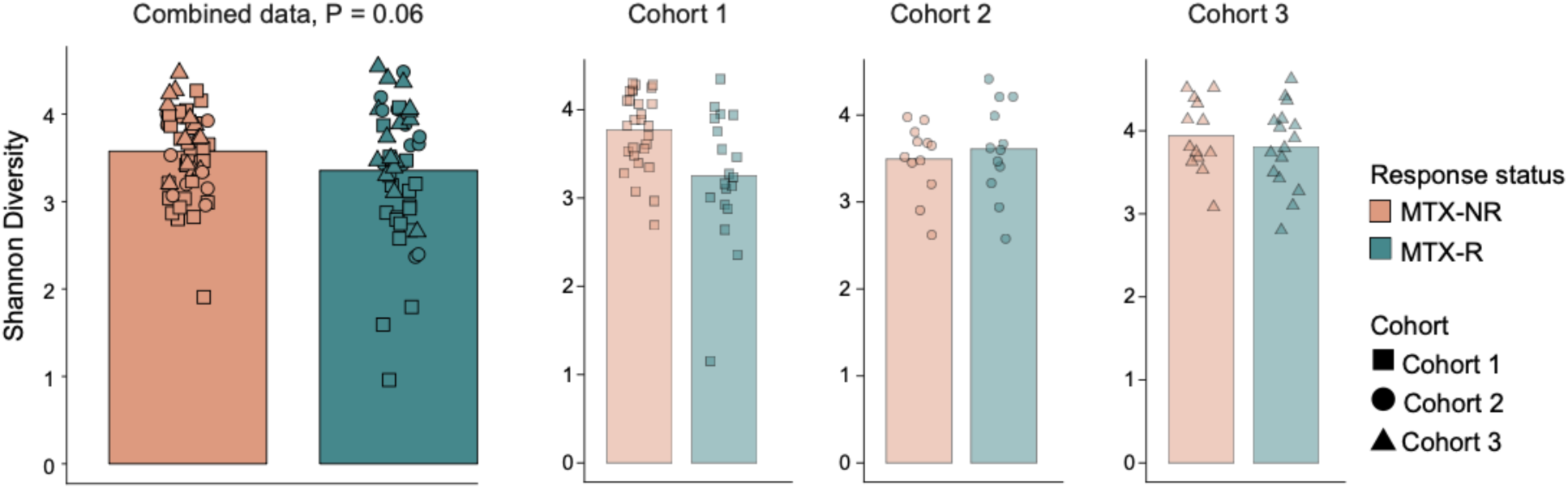
Pretreatment gut microbiome alpha diversity does not differ by future MTX response. Shannon diversity of strains shown across all cohorts and within each individual cohort. Bars represent group means, with individual samples overlaid as points. Statistical comparisons were performed using the Welch’s t-test with p values indicated.

**Supplementary Figure 5.**
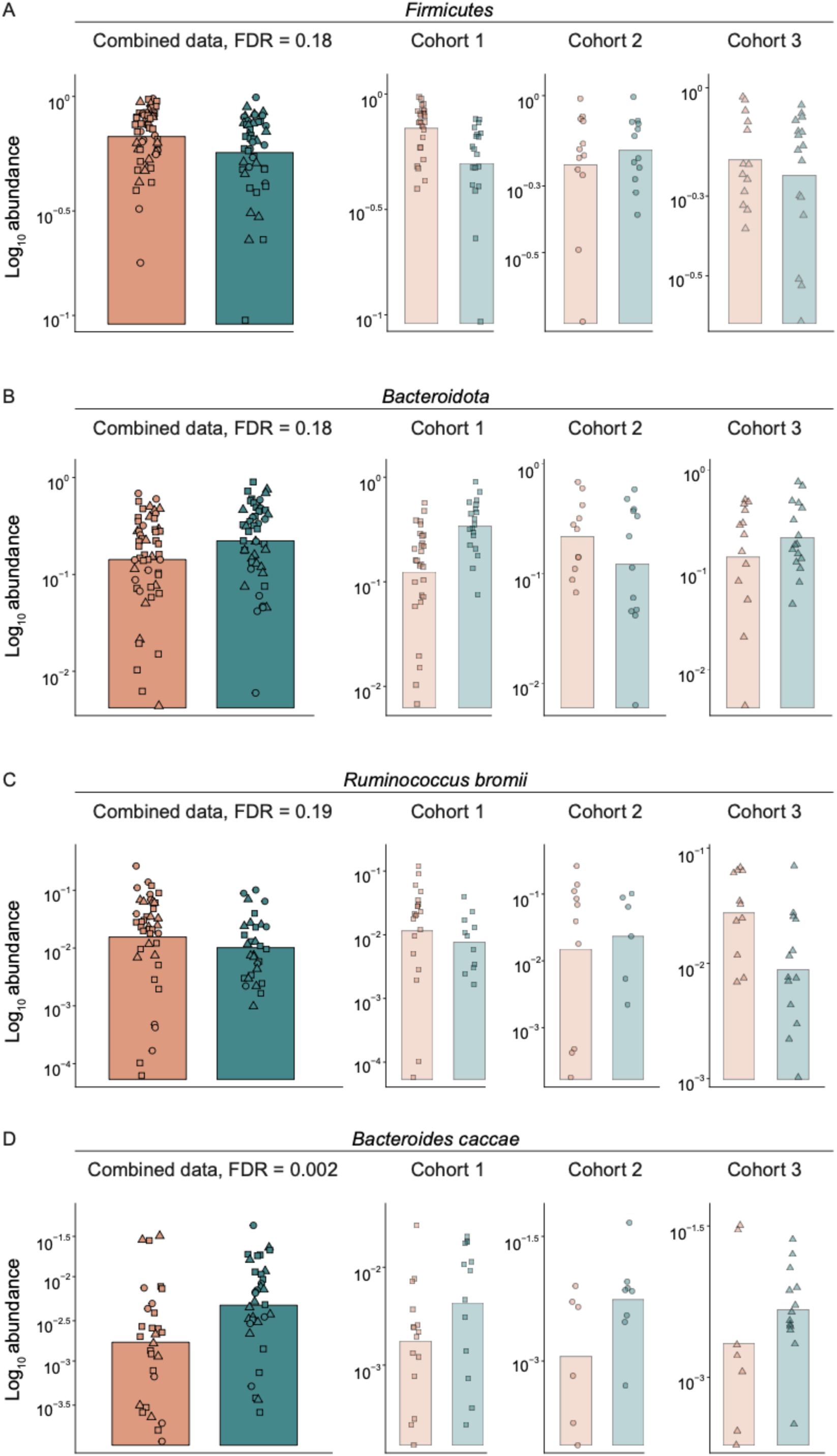
MTX-NR exhibits lower abundance of *Bacteroidota* and higher abundance of *Firmicutes*. (**A**) Log scaled abundance of *Bacteroidota* (**B**) *Firmicutes* (**C**) *Ruminococcus bromii*, and (**D**) *Bacteroides caccae* in MTX-R and MTX-NR across all cohorts and within each cohort. Bars represent group means, with individual samples overlaid as points. Linear regression was performed adjusting for age, sex, and cohort. FDR values as indicated.

**Supplementary Figure 6.**
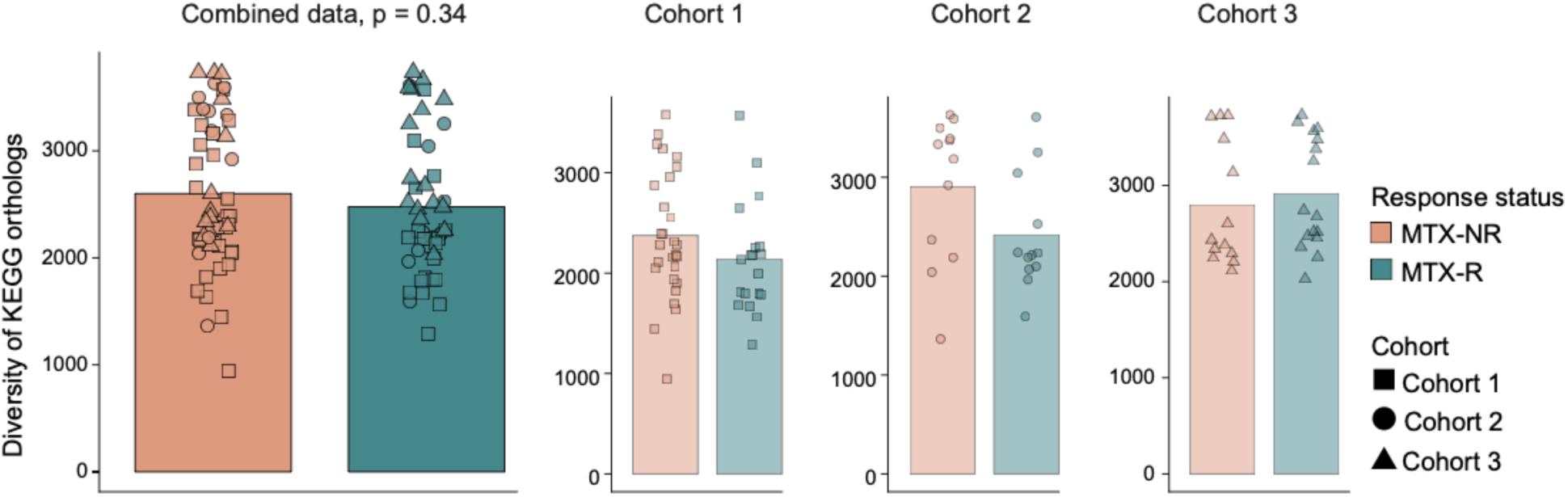
The number of distinct KEGG orthologs (alpha diversity) in MTX-R and MTX-NR did not differ by treatment response. Richness of KEGG orthologs in MTX-R and MTX-NR is shown across all cohorts and within each individual cohort. Bars represent group means, with individual samples overlaid as points. Statistical comparisons were performed using the Wilcoxon rank-sum test, with *p*-values indicated.

**Supplementary Figure 7.**
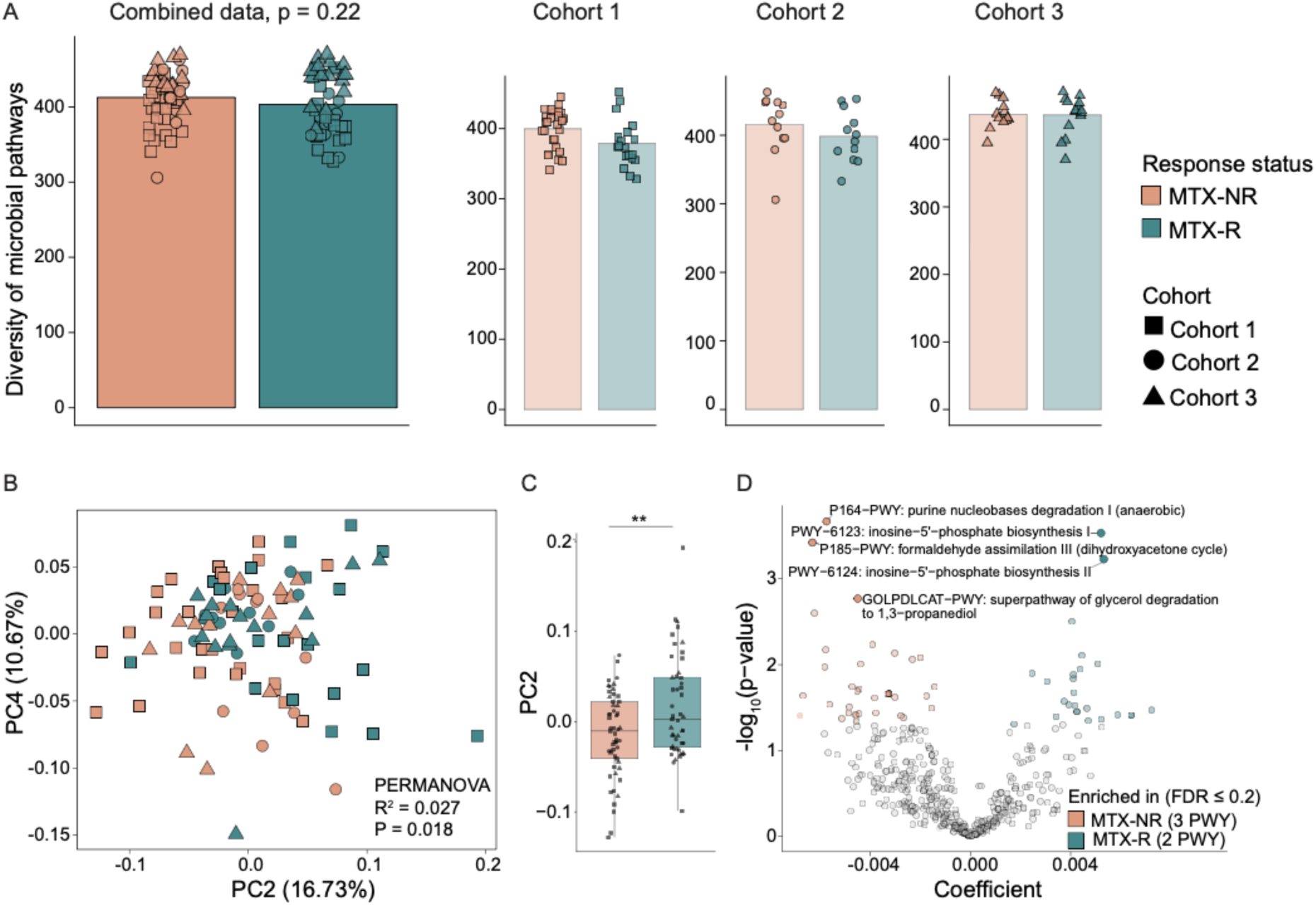
The pretreatment composition of gut microbial pathways differs between MTX-R and MTX-NR in the NORA cohorts. (**A**) Richness of gut microbial pathways in MTX-R and MTX-NR is shown across all cohorts and within each individual cohort. Bars represent group means, with individual samples overlaid as points. Statistical comparisons were performed using the Wilcoxon rank-sum test, with *p*-values indicated. (**B**) PCoA of gut microbial pathway composition based on Bray-Curtis distance (PERMANOVA, R² = 0.027, *p* = 0.018). Box plots along PCoA2 and PCoA4 show the distribution of sample values along each axis, and Welch’s t-test p-values are reported. (**C**) Boxplots representing score of PCoA component 2. Significance is indicated as follows: \*\**p* < 0.01 (Wilcoxon rank sum test) (**D**) Differentially abundant gut microbial pathways between MTX-R and MTX-NR groups. Linear regression meta-analysis was performed using MMUPHin, adjusting for age, sex, and cohort. Results are shown as a volcano plot, where colored points represent significantly different pathways (solid points, FDR ≤ 0.2; transparent points, unadjusted p < 0.05).

**Supplementary Figure 8.**
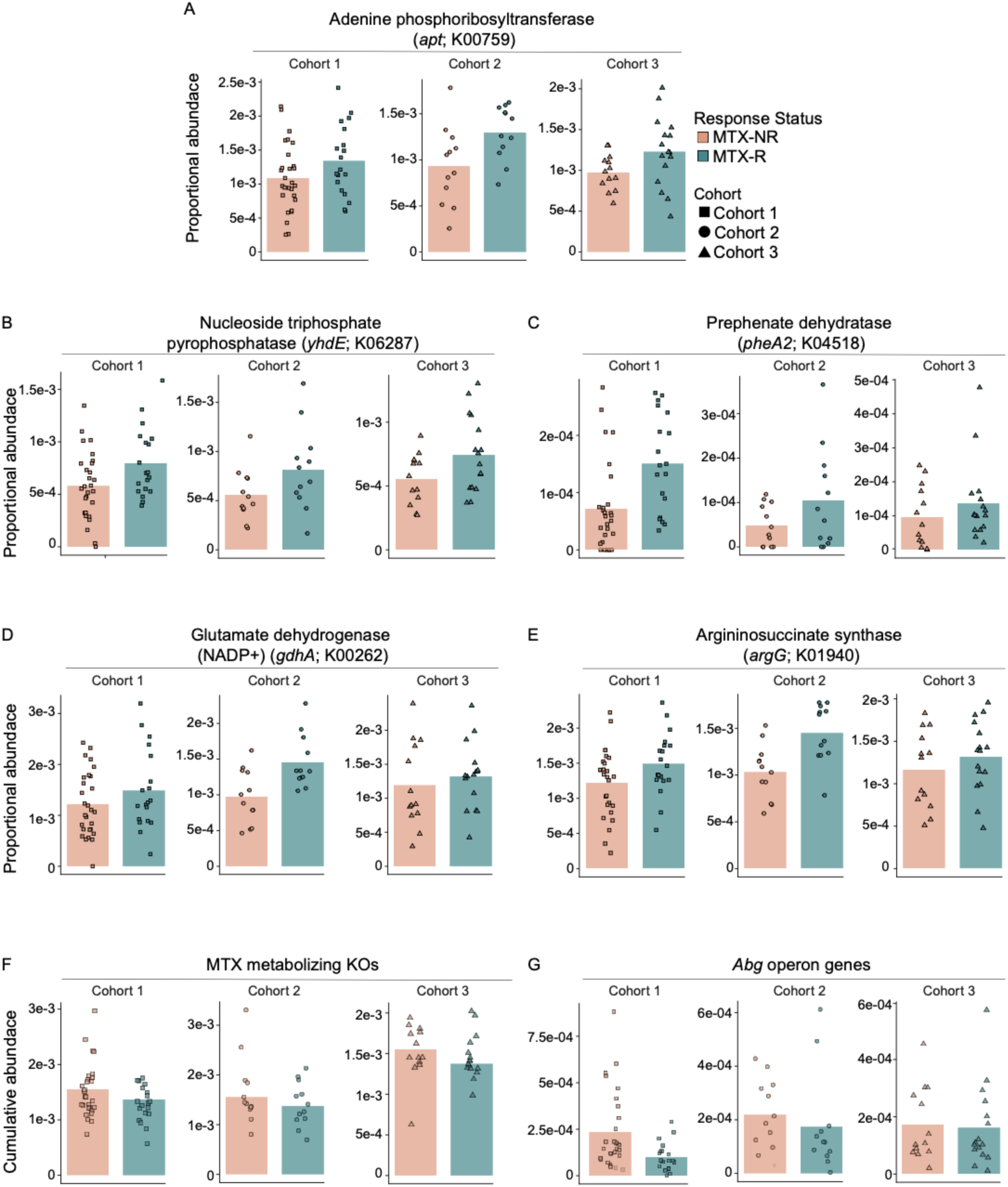
Cohortwise abundance of KOs involved in Nucleotide metabolism and amino acid synthesis and MTX metabolism in MTX-R and MTX-NR. Abundance of **(A)** Adenine phosphoribosyltransferase (*apt*; K00759), **(B)** nucleotide triphosphate pyrophosphatase (*yhdE*; K06287), **(C)** prephenate dehydratase (*pheA2*; K04518), **(D)** glutamate dehydrogenase (*gdhA*; K00262), and **(E)** argininosuccinate synthase (*argG*; K01940). Data are presented as proportional abundance. Cumulative abundance of (**F**) 9 KEGG orthologs predicted to metabolize MTX^19^ (**Supplementary Table 9**) and of **(G)** *abg* operon genes (*abgA*, *abgB*, *abgT*) (**Supplementary Table 10**) in each cohort. Data are presented as proportional abundance values. Bars represent group means, with individual samples overlaid as points.

**Supplementary Figure 9.**
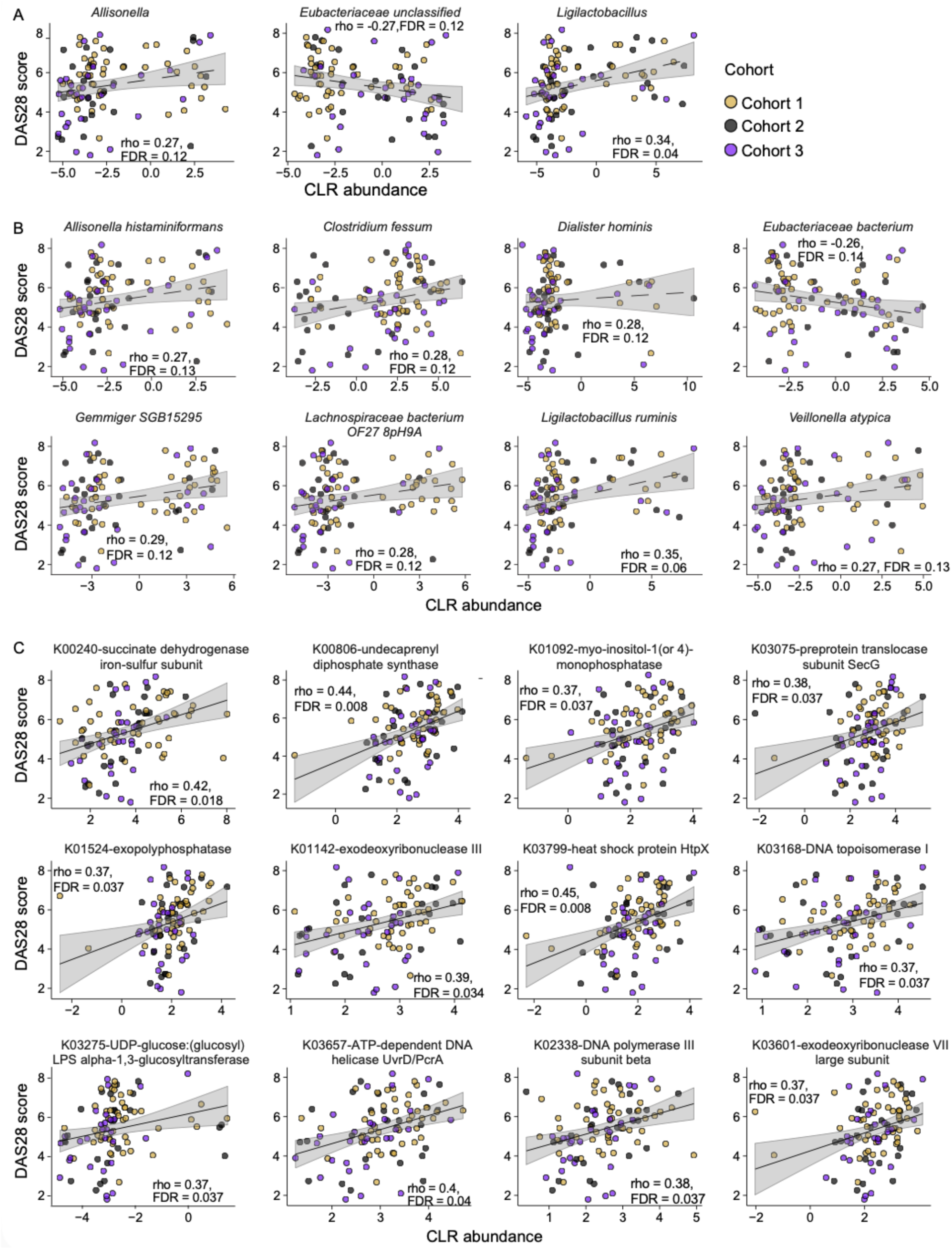
Baseline DAS28 scores are associated with bacterial genera, species and gene families in NORA subjects. Spearman’s rank correlation was performed between DAS28 scores and the CLR abundance of **(A)** bacterial genera, **(B)** species and (**C**) KEGG orthologs. The y-axis represents DAS28 scores and x-axis represent microbial taxa or KEGG features.

**Supplementary Figure 10.**
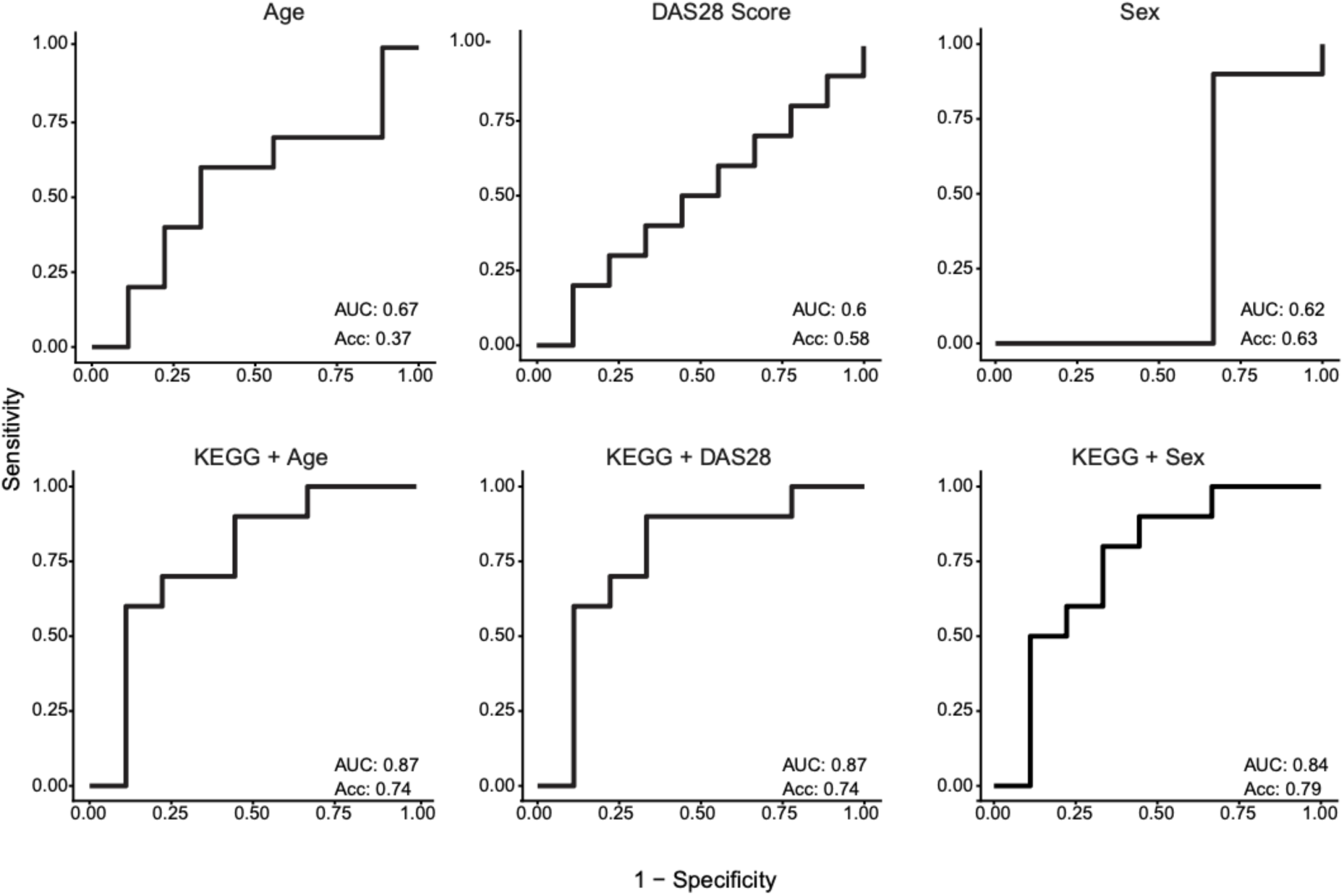
Clinical parameters poorly predict response to MTX treatment. Random forest models were built using clinical variables (DAS28, age, sex), either individually in combination with Boruta-selected gene orthologs (KEGG). Models were built using 80% of samples, and the remaining 20% were used to test the model. The area under the curve (AUC) obtained with each model is shown.

